# Identification of a mitochondrial targeting sequence in cathepsin D and its localization in mitochondria

**DOI:** 10.1101/2023.01.23.524639

**Authors:** Naoki Ikari, Hirofumi Arakawa

**Author notes:** Correspondence should be addressed to: Hirofumi Arakawa, M.D., Ph.D., Division of Cancer Biology, National Cancer Center Research Institute, 5-1-1 Tsukiji, Chuo-ku, Tokyo 104-0045, JAPAN.

## Abstract

Cathepsin D (CTSD) is a major lysosomal protease harboring an N-terminal signal peptide (amino acids 1–20) to enable vesicular transport from endoplasmic reticulum to lysosomes. Here, we report the possibility of a mitochondrial targeting sequence and mitochondrial localization of CTSD in cells. Live-cell imaging analysis with enhanced green fluorescent protein (EGFP)-CTSD indicated that CTSD localizes to mitochondria. CTSD amino acids 21–35 are responsible for its mitochondrial localization, which exhibit typical features of mitochondrial targeting sequences, and are evolutionarily conserved. A proteinase K protection assay and sucrose gradient analysis showed that a small population of endogenous CTSD molecules exists in mitochondria. These results suggest that CTSD is a dual-targeted protein that may localize in both lysosomes and mitochondria.

## Introduction

CTSD is an aspartic lysosomal protease, dysfunction of which causes the most severe congenital form of neuronal ceroid lipofuscinosis (NCL), characterized by neurodegeneration and accumulation of ceroid lipofuscin containing the mitochondrial F1F0 ATPase [1, 2]. To date, the process of lysosomal localization of CTSD has been characterized as follows [3, 4]. Human CTSD, comprising 412 amino acids (pre-pro-CTSD), is translocated into endoplasmic reticulum (ER) via the ER signal peptide (amino acids 1-20), which is removed during translocation to yield pro-CTSD (392 amino acids, 52 kDa) [3]. After undergoing N-linked glycosylation (amino acids 134 and 263) in ER and subsequent modifications in Golgi [5, 6], pro-CTSD is transported to the endolysosomal compartment via mannose-6-phosphate (M6P)-dependent or independent pathways [4, 7]. Upon entering the acidic endolysosomal compartment, the N-terminal propeptide (amino acids 21-64) is proteolytically processed to a 48-kDa single-chain intermediate form. Finally, the intermediate form is processed into the mature enzymatic form, comprising heavy (34 kDa) and light (14 kDa) chains linked by non-covalent interactions, being cleaved by cathepsins B and L [8] and progranulin [9].

After lysosomal localization, CTSD is reportedly secreted to the extracellular space in breast cancer cells [10]. Other studies have found that CTSD is also released from lysosomes to the cytosol under stresses such as acetic acid [11] and staurosporine [12].

However, several studies have detected CTSD in mitochondria. i) The fruit fly ortholog of CTSD was detected in mitochondrial extracts using IEF/SDS-PAGE [13]. ii) The yeast ortholog of CTSD was detected in a 44.5-kDa form during proteomic analyses of mitochondria [14]. iii) The mouse ortholog of CTSD was detected during O-GlcNAcylome analysis of isolated mitochondria [15]. iv) Human CTSD was detected in the mitochondrial fraction of colon cancer cells with Mieap/SPATA18 expression [16], and v) human CTSD was detected in the mitochondrial fraction of gastric cancer cells in a 48-kDa form during hypoxia experiments [17]. These reports of CTSD in mitochondria cannot be explained by protein transport from the ER to lysosomes. Nevertheless, no mechanism for this phenomenon has been suggested.

In this study, to further explore subcellular localization of CTSD, we performed live-cell imaging analysis of EGFP-CTSD using various cell lines. We confirmed that EGFP-CTSD is localized not only in lysosomes, but also in mitochondria. Furthermore, sequence analysis, live-cell imaging analysis using deletion mutants of EGFP-CTSD in live cells, a proteinase K protection assay and sucrose gradient analysis for endogenous CTSD, and conventional immunofluorescence (IF) analysis suggest that there is a mitochondrial targeting sequence (MTS) in CTSD, and that a subpopulation of physiological CTSD may localize in mitochondria via its MTS. On the basis of these results, we conclude that CTSD localizes in mitochondria via its MTS.

## Results

### EGFP-CTSD localizes in mitochondria

To examine subcellular localization of CTSD in cells, we first constructed a plasmid to express C-terminal EGFP-tagged CTSD (EGFP-CTSD) and performed live-cell imaging of EGFP-CTSD using three cancer cell lines (HeLa, SK-BR-3, and HepG2 cells), and a non-cancer cell line (293 cells). Lysosomal localization of EGFP-CTSD was observed in all four cell lines (Hela, 45.9±7.3%; SK-BR-3, 39.7±1.4%; HepG2, 60.1±4.9%; 293, 58.7±2.6%) (Fig. 1A –D and 1I). Surprisingly, EGFP-CTSD signals were also detected in mitochondria of all four cell lines (HeLa, 42.6±6.8%; SK-BR-3, 78.7±9.5%; HepG2, 42.7±6.1%; 293, 38.3±2.2%) (Fig. 1E – H and 1J). Interestingly, we frequently observed that mitochondrial and lysosomal localization of EGFP-CTSD were mutually exclusive in individual cells. Thus, these results suggest that regardless of cellular profiles, CTSD does localize in mitochondria.

**Fig. 1.**
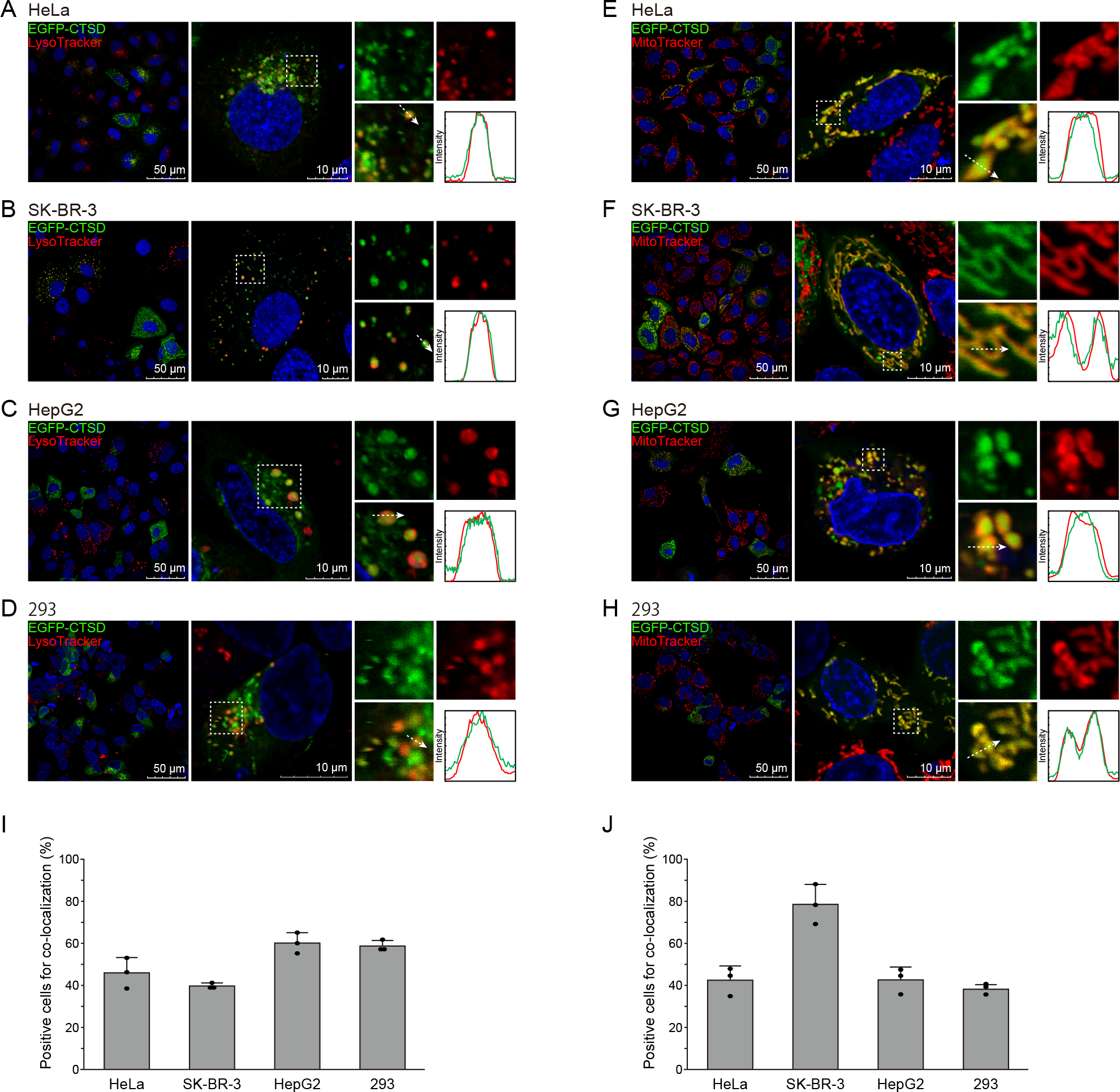
Mitochondrial localization of EGFP-CTSD. **(A–J)** Live-cell imaging showing the subcellular localization of EGFP-CTSD expressed in HeLa **(A and E)**, SK-BR-3 **(B and F)**, HepG2 **(C and G)**, and 293 cells **(D and H)** compared with a lysosome probe, LysoTracker Red **(A–D)** and a mitochondria probe, MitoTracker Red **(E–H)**. **(I and J)** Co-localization rates of EGFP-CTSD with LysoTracker Red **(I)** and MitoTracker Red **(J)** in HeLa, SK-BR-3, HepG2, and 293 cells, as in **(A–J)**. n = 103 – 192 cells from three independent experiments. Data shown are means ±SD. Raw data are provided in Supplementary Table S2.

### CTSD has an evolutionary conserved MTS

To identify a potential MTS in the sequence of CTSD, we performed sequence analyses. First, we used DeepLoc-1.0 and DeepLoc-2.0, because DeepLoc-1.0 can depict a hierarchical tree likelihood, mimicking a subcellular sorting pathway [21], and DeepLoc-2.0 is improved for multi-localization prediction [22]. Consistent with evidence that the N-terminal 1–20 amino acids of CTSD is an ER signal peptide for vesicular transport from the ER to lysosomes [3], DeepLoc-1.0 sorted CTSD toward lysosomes (Fig. 2A) [21], and DeepLoc-2.0 predicted that CTSD has an ER signal peptide and localizes in lysosomes and vacuoles (Fig. 2B) [22].

**Fig. 2.**
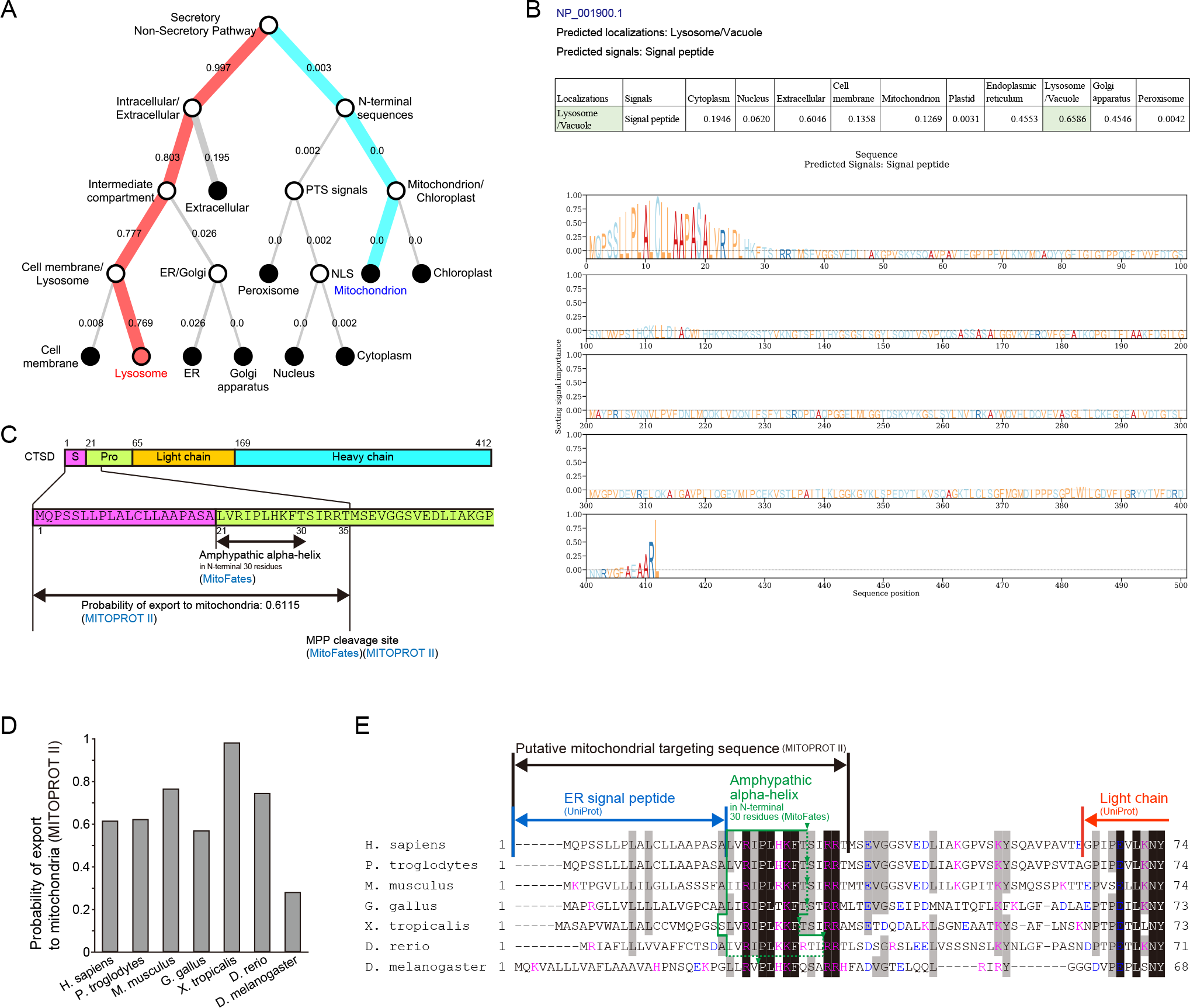
CTSD has a predicted MTS containing an evolutionarily conserved region (amino acids 21-35). **(A)** Prediction of subcellular localization of CTSD by DeepLoc-1.0, depicted in a hierarchical tree diagram. **(B)** Prediction of subcellular localization of CTSD by DeepLoc-2.0. The predicted localization is highlighted in green (upper panel). Regions in the sequence responsible for the predicted localization are shown as high values in the Logo-like plot (lower panel). **(C)** Prediction of mitochondrial localization and the MTS of CTSD by MITOPROT II and MitoFates. **(D)** Prediction of mitochondrial localization of CTSD orthologs in representative eukaryotes by MITOPROT II. **(E)** Multiple sequence alignment for CTSD orthologs in representative eukaryotes. Black and gray boxes indicate 100% and 70% identical residues among eukaryotes, respectively. Pink letters indicate positively charged residues. Blue letters indicate negatively charged residues.

On the other hand, when we used MITOPROT II, it predicted amino acids 1–35 of CTSD as a putative MTS, and probable transport of CTSD to mitochondria as 0.6115 [24] (Fig. 2C). It has been suggested that dual-localized proteins often exhibit lower MITOPROT II scores, compared to typical mitochondrial proteins. The median value of dual-localized proteins is 0.603 [25]. MITOPROT II scores for several CTSD orthologs also revealed this feature (Fig. 2D).

In addition, when we used MitoFates [23], it predicted at least 21–30 amino acids of CTSD as an amphipathic alpha-helix (Fig. 2C). Thus far, the 3D structure of amino acids 21–35 has not been determined, according to the Protein Data Bank (PDB) (https://www.rcsb.org/, 6th May 2022) [26]. When we performed multiple sequence alignment for CTSD orthologs in representative eukaryotes, we found that amino acids 21–35 are evolutionarily conserved, showing enrichment of positively charged residues and a paucity of acidic residues (Fig. 2E). We also found that there is a putative mitochondrial protein peptidase (MPP) cleavage site at amino acids 35/36 of CTSD (Fig. 2C) [23, 24]. These are all characteristic features of MTS [24, 27, 28].

### The predicted MTS of CTSD is required for mitochondrial localization of EGFP-CTSD

To verify the putative MTS predicted by sequence analyses for CTSD, we examined cells expressing EGFP-CTSD-full (Full-length) and four deletion-mutant forms, EGFP-CTSD Δ2–20 (without the ER signal peptide), Δ2–35 (without the ER signal peptide and the putative MTS predicted by MITOPROT II [24]), Δ2–64 (without the ER signal peptide, the putative MTS, and propeptide), and Δ21–35 (without the putative MTS) (Fig. 3A).

**Fig. 3.**
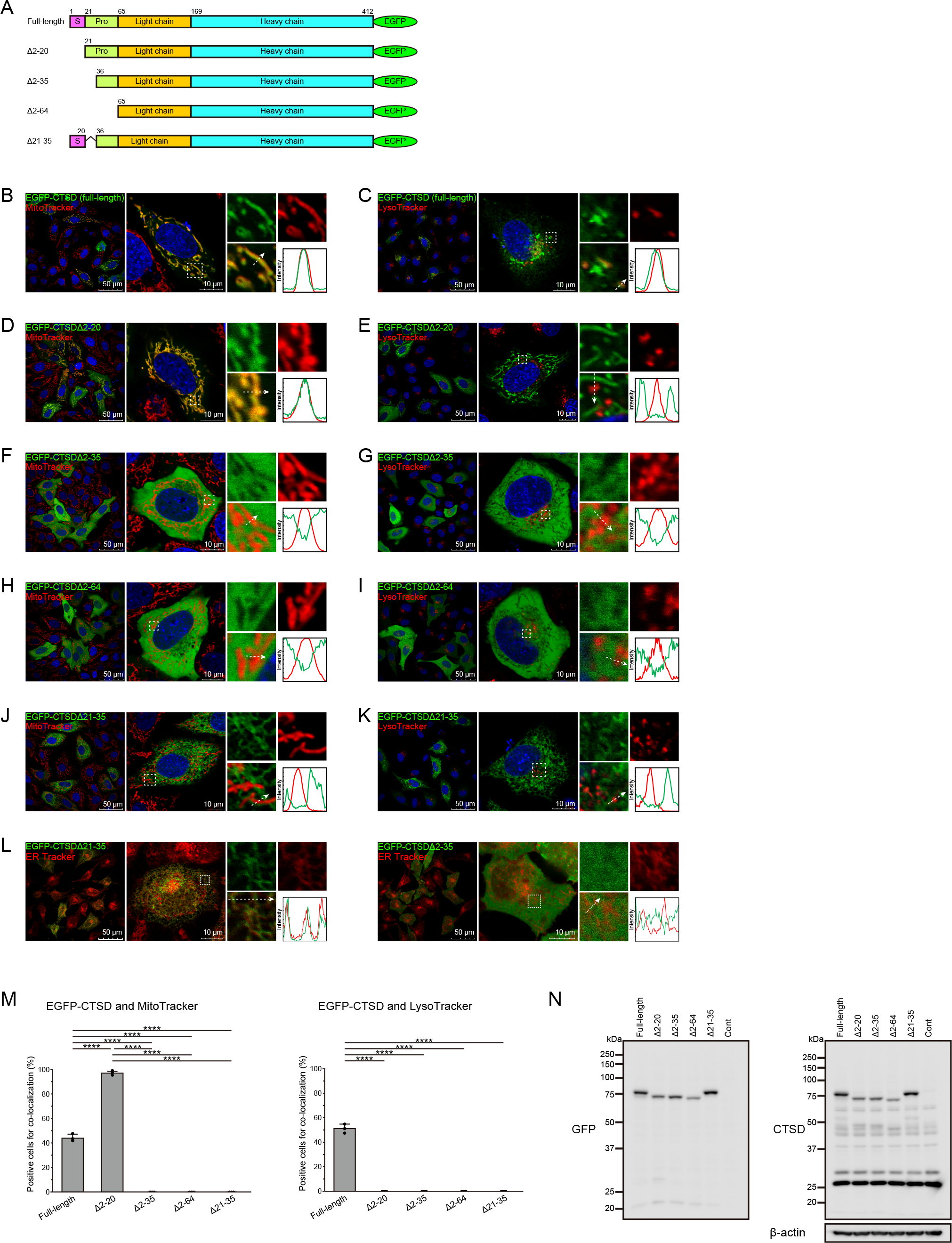
CTSD amino acids 21-35 comprise a bona-fide MTS, responsible for mitochondrial localization. **(A)** A schematic of EGFP-CTSD and deletion mutants (Δ2–20, Δ2–35, Δ2–64, and Δ21–35). Numbers indicate amino acids. **(B–K)** Live-cell imaging showing subcellular localization of EGFP-CTSD full-length **(B and C)**, Δ2–20 **(D and E)**, Δ2–35 **(F and G)**, Δ2–64 **(H and I)**, and Δ21–35 **(J and K)** expressed in HeLa cells, compared with a mitochondrial probe, MitoTracker Red **(B, D, F, H, J)** and a lysosomal probe, LysoTracker Red **(C, E, G, I, K)**. **(L)** Live-cell imaging showing subcellular localization of EGFP-CTSDΔ21–35 (left) and Δ2–35 (right) compared with an endoplasmic reticulum (ER) probe, ER Tracker Blue-White DPX in HeLa cells. **(M)** Co-localization rates of EGFP-CTSD full-length, Δ2–20, Δ2–35, Δ2–64, and Δ21–35 expressed in HeLa cells with MitoTracker Red (left) and LysoTracker Red (right). n = 104 – 165 cells from three independent experiments. Data shown are means ±SD. Asterisks indicate significance levels as follows: ****, p < 0.0001. **(N)** Western blot to evaluate cleavage of EGFP-CTSD expressed in HeLa cells, in comparison with molecular weights of deletion mutants (Δ2–20, Δ2–35, Δ2–64, and Δ21–35) expressed in HeLa cells. Raw data are provided in Supplementary Table S3. Full-gel images are provided in Supplementary Fig. S1.

Reproducibly as in Fig. 1E, cells expressing full-length EGFP-CTSD showed dual localization of CTSD in mitochondria and lysosomes (43.9±3.3% and 51.1±1.3% of expressing cells, respectively) (Fig. 3B, C, and M). Cells expressing Δ2–20, which lacks only the ER signal peptide, showed more intensive mitochondrial localization (96.8±1.5% of expressing cells) and loss of lysosomal localization (Fig. 3D, E, and M) of CTSD. Mitochondrial localization of CTSD was lost in cells expressing Δ2–35, Δ2–64, and Δ21–35, all of which lack amino acids 21–35 of CTSD (Fig. 3F–K, and 3M). Among them, Δ2–35 and Δ2–64 lacking the ER signal peptide were distributed all through the cytoplasm (Fig. 3F–I, and 3M), and Δ21–35 cells, retaining the ER signal peptide, were distributed to ER, but were not delivered to lysosomes (Fig. 3J–L, and 3M). These data indicate that the MTS of CTSD is limited to amino acids 21–35 of CTSD, and potentially competes with the ER signal peptide.

We performed western blot (WB) analysis for cells expressing EGFP-CTSD and deletion mutants. As shown in Fig. 3N, there were no bands equivalent to or smaller than Δ2–35 in cells expressing EGFP-CTSD full-length, suggesting that at least EGFP-tagged CTSD is not cleaved at the putative MPP cleavage site (amino acids 35/36) predicted by sequence analyses (Fig. 2C). One caveat here is that there were also no bands similar to the band of Δ2–64 (corresponding to the intermediate form) or smaller than Δ2–64 (corresponding to the mature form) in cells expressing EGFP-CTSD full-length. This implies that EGFP-tagged CTSD may be impaired in processing for transport from ER to lysosomes (Fig. 3N). In summary, CTSD amino acids 21–35 comprise a bona fide MTS, and at least EGFP-tagged CTSD is not cleaved at the putative MPP cleavage site (amino acids 35/36) predicted by sequence analyses.

### A small population of endogenous CTSD exists in mitochondria

To examine whether endogenous CTSD exists in mitochondria, we performed a proteinase K protection assay with HeLa cells. In advance, we confirmed WB bands of endogenous CTSD variants using CTSD-knockdown (KD) cells and two anti-CTSD antibodies (CTD-19, mouse monoclonal and sc-6486 goat polyclonal). Multiple bands proved specific for endogenous CTSD expression, which was detected in 43~48-kDa (precursor form) and 30-kDa (active form) regions (Fig. 4A).

**Fig. 4.**
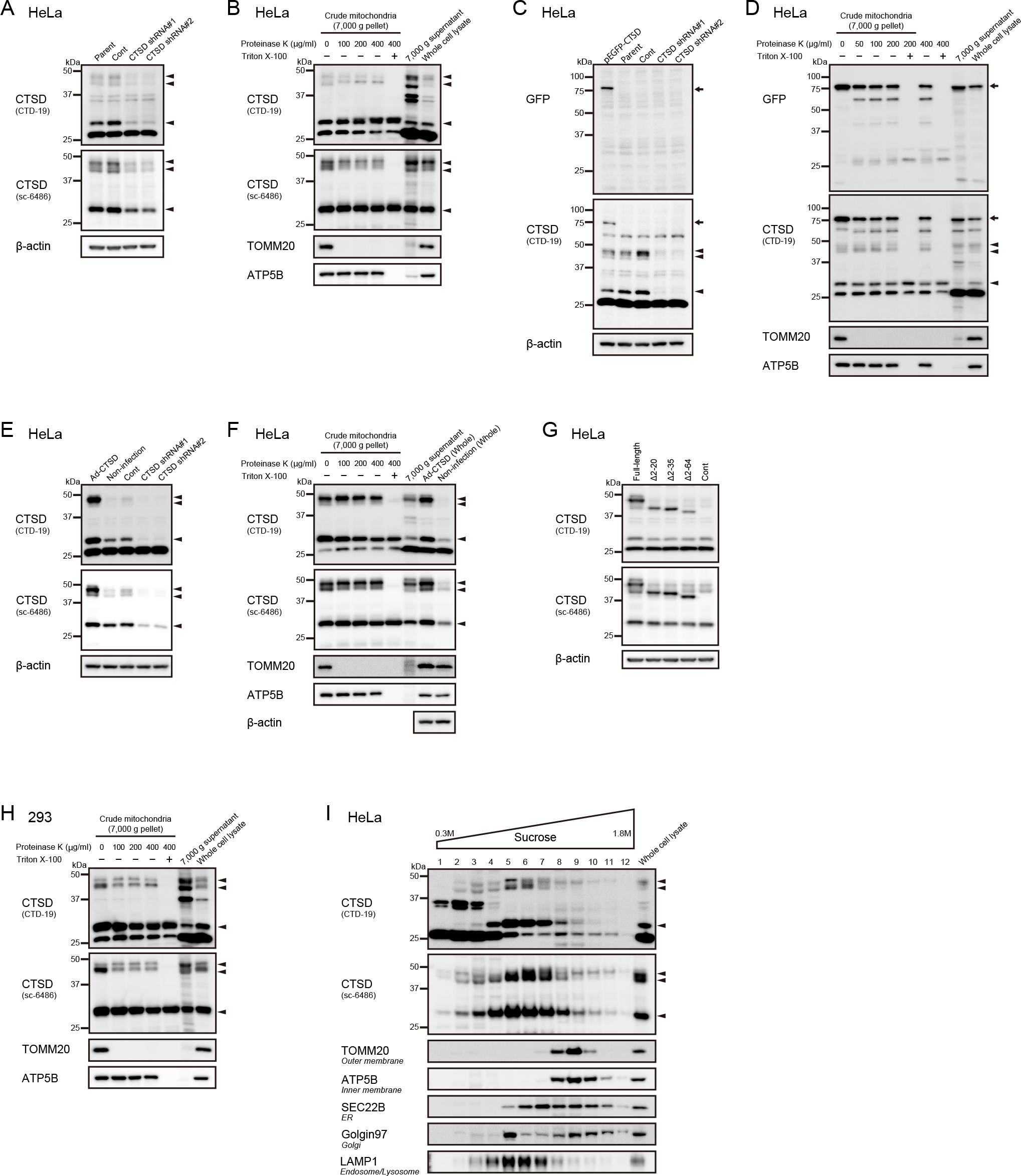
A small amount of endogenous CTSD is localized in mitochondria. **(A)** Verification of specific bands of endogenous CTSD in Western blots. The HeLa parent, control-KD, and two CTSD-KD cells expressing different short-hairpin RNAs against the CTSD sequence (shRNA#1 and shRNA#2) are compared. Two anti-CTSD antibodies (CTD-19, mouse monoclonal; sc-6486 goat polyclonal) were used for accuracy. Arrowheads indicate specific bands of endogenous CTSD. **(B)** Proteinase K protection assay for HeLa cells. The crude mitochondrial fraction (7,000 g pellet) isolated from HeLa cells was treated with increasing concentrations of proteinase K in the presence and absence of Triton X-100, followed by Western blotting. The remaining 7,000 g supernatant and whole cell lysates were applied to adjacent lanes for comparison. Arrowheads indicate specific bands of endogenous CTSD verified in **(A)**. **(C and D)** For HeLa cells expressing EGFP-CTSD, verification of bands of EGFP-CTSD and endogenous CTSD **(C)** and proteinase K protection assay **(D)** were performed as in **(A and B)**. Arrows and arrowheads indicate bands of EGFP-CTSD and endogenous CTSD, respectively. **(E and F)** For HeLa cells expressing exogenous CTSD, verification of bands of CTSD **(E)** and proteinase K protection assay **(F)** were performed as in **(A and B)**. Arrowheads indicate bands of CTSD. **(G)** Comparison of ~50kDa bands that contained proteinase K-inaccessible CTSD in **(B)** with bands of CTSD-deletion mutants (Δ2–20, Δ2–35, Δ2–64, and Δ21–35) in Western blots. **(H)** For 293 cells, the proteinase K protection assay was performed as in **(B)**. **(I)** Sucrose gradient analysis for HeLa cells. HeLa cell extract was fractionated by sucrose density gradient centrifugation. Fractions were collected and subjected to Western blot analysis, using antibodies against indicated proteins. Full-gel images are provided in Supplementary Fig. S2–S7.

A crude mitochondrial fraction isolated from Hela cells was treated with proteinase K in the presence and absence of Triton X-100. TOMM20 (an outer mitochondrial membrane protein) and ATP5B (the matrix face of the inner mitochondrial membrane protein) were used as mitochondrial protein controls. A subpopulation comprising bands of 43~48-kDa were protected by proteinase K digestion, suggesting that a small proportion of endogenous CTSD may exist in mitochondria (Fig. 4B). On the other hand, the 30-kDa band was retained upon proteinase K digestion, even in the presence of Triton X-100. This suggests that this band is resistant to proteinase K (Fig. 4B), so it could not be evaluated using a proteinase K protection assay.

We also examined whether exogenous EGFP-CTSD or exogenous CTSD is protected from proteinase K digestion. Both EGFP-CTSD and exogenous CTSD were much more clearly concentrated in the mitochondrial fraction and protected by proteinase K digestion, compared to endogenous CTSD (Fig. 4C–F).

To examine whether mitochondrial forms of endogenous CTSD contain the putative MTS (amino acids 21-35), we compared multiple bands of endogenous CTSD, which were protected from proteinase K digestion, with each band of full-length and Δ2–20, Δ2–35, and Δ2–64 mutants of exogenous CTSD in WB. Mitochondrial forms of endogenous CTSD are likely consistent with full-length and Δ2–20, but larger than Δ2–35 and Δ2–64 (Fig. 4G). This result is consistent with our finding in the imaging analysis with their mutants (Fig. 3) that the MTS of CTSD is amino acids 21– 35 of CTSD. In addition, consistent with the results of EGFP-CTSD, mitochondrial forms of endogenous CTSD may not be cleaved at the putative MPP cleavage site (amino acids 35/36).

To validate the mitochondrial forms of endogenous CTSD in normal cells, we performed a proteinase K protection assay with 293 cells. Similar results were obtained with 293 cells, suggesting that mitochondrial localization of endogenous CTSD may be common to all cell types (Fig. 4H).

To further confirm the possible existence of endogenous CTSD in mitochondria, we performed sucrose gradient analysis with HeLa cell extract. A small population of endogenous CTSD was detected in the mitochondrial fraction (lanes 8-10), indicated by bands of TOMM20 and ATP5B (Fig. 4I). This result also supports the notion that a small part of endogenous CTSD exists in mitochondria.

### Immunostaining experiments with fixed cells may fail to detect mitochondrial CTSD protein

Thus far, to our knowledge, endogenous CTSD has never been detected in immunofluorescence (IF) experiments using fixed cells. Therefore, to examine whether mitochondrial CTSD is stable in fixed cells, which are usually used in IF experiments, we performed fixed-cell imaging with cells expressing EGFP-CTSD, comparing it with imaging of live-cells (Fig. 5A).

**Fig. 5.**
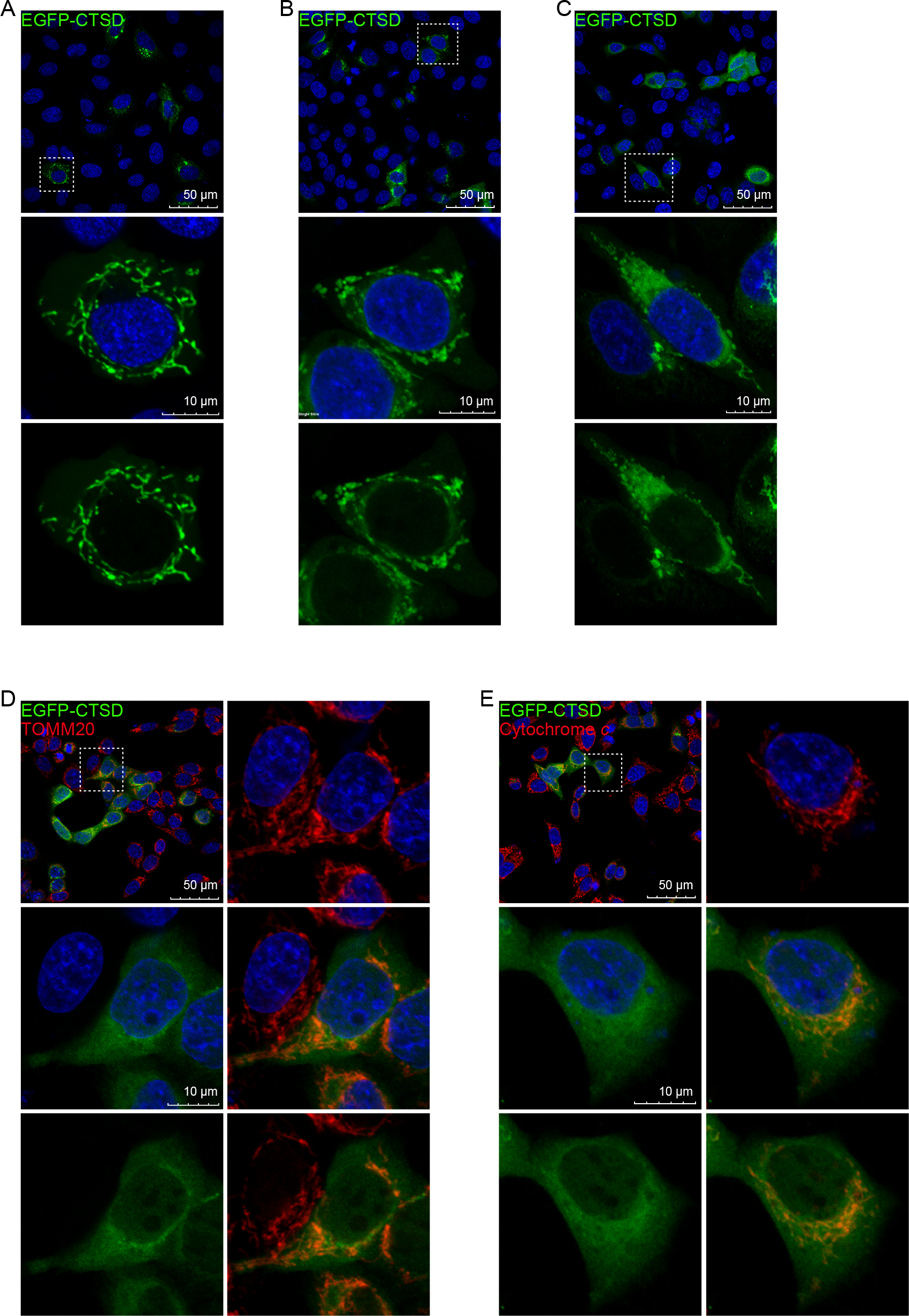
Mitochondrial EGFP-CTSD cannot be detected by IF with fixed cells. **(A–E)** Live-cell imaging **(A)** and fixed-cell imaging **(B–E)** of HeLa cells expressing EGFP-CTSD with the following procedures. Fixation with 4% PFA for 15 min **(B)**, with permeabilization in 0.1% Triton X-100 for 5 min **(C)**, and immunoreaction **(D and E)** using anti-TOMM20 antibody **(D)** and anti-cytochrome *c* antibody **(E)**.

Compared to live imaging data (Fig. 5A), localization of EGFP-CTSD seemed to be prevented by a gentle fixation procedure (4% PFA 15 min) without a permeabilization procedure (Fig. 5B). This implies that, even when we skipped the cell permeabilization procedure, subsequent general procedures required for IF may cause diffusion of mitochondrial EGFP-CTSD in the cell. When we performed a permeabilization procedure, which is essential to detect proteins in mitochondria surrounded by a double-membrane (0.1% Triton X-100 5 min), mitochondrial localization of EGFP-CTSD was more corrupted (Fig. 5C) or completely lost with immunostaining procedures, though staining of TOMM20 and cytochrome *c* was still maintained (Fig. 5D and 5E). These results suggest that immunostaining experiments with fixed cells may fail to detect mitochondrial localization of endogenous CTSD.

## Discussion

Here, we show that, upon live-cell imaging, EGFP-CTSD (known as lysosomal protease) localizes in mitochondria. Amino acids 21–35 of CTSD were responsible for mitochondrial localization of CTSD, which presented typical features of an MTS and were evolutionarily conserved. Furthermore, a subpopulation of endogenous CTSD was protected from proteinase K, and detected in mitochondrial fractions by sucrose gradient analysis, indicating that a small proportion of endogenous CTSD could be localized in mitochondria. Molecular weights of mitochondrial forms of CTSD were higher than those of CTSD mutants without the MTS in WB, which suggests that mitochondrial forms are not cleavable at a putative MPP site in mitochondria. We also found that mitochondrial CTSD was unstable in fixed cells. That could explain why it has not been detected in past IF experiments.

In the present study, we identified a previously unknown mitochondrial localization of CTSD, one of the most investigated lysosomal proteases [3, 4, 29]. Although mitochondrial targeting mechanisms for subcellular proteins are conservative [30], the mitochondrial localization of CTSD has been overlooked by more than 6,400 studies in PubMed until this discovery. Since mitochondrial localization of CTSD has been confirmed in various cell types, and the MTS is conserved across all species examined, this phenomenon may be universal. In addition, since CTSD is associated with various common diseases including neurodegenerative diseases [31], cancer [32] [33], and atherosclerosis [34], our finding may stimulate a lot of interest and discussion regarding associated pathophysiology and possible therapeutic strategies.

Among 798 proteins that fulfill criteria for mitochondrial localization, including genome-wide experimental screens [35], 316 proteins are potentially dual-targeted, with a second location prone to be concealed by uneven distribution [35, 36]. However, in our study, mitochondrial CTSD was clearly visualized by live-cell imaging analysis of over-expressed EGFP-CTSD. Thus, live-cell imaging analysis of over-expressed proteins with optimized fluorescent tags could be a powerful tool for detecting potential dual-targeted proteins to overcome “eclipsed distributions”, probably comprising a considerable number of proteins [35, 36].

Why has CTSD never been detected in past imaging analyses? We suggest several possible reasons. (1) Live-cell imaging analysis of over-expressed EGFP-CTSD may have not been performed in previous studies. (2) Since the amount of mitochondrial endogenous CTSD is very small, these forms may have been overlooked in past biochemical analyses, such as WB. The mitochondrial volume is much larger than the lysosomal volume in cells, suggesting that further dilution of a small amount of endogenous CTSD may have made detection difficult. (3) Immunostaining experiments with fixed cells fail to detect mitochondrial CTSD. In this study, we demonstrated the fragility of mitochondrial CTSD proteins in IF experiments by using EGFP-CTSD and fixed cells. This feature of mitochondrial CTSD in IF experiments may have made it additionally difficult to detect mitochondrial CTSD previously.

What is the rationale for mitochondrial localization of CTSD? CTSD requires an acidic pH to be proteolytically active [29]. In this regard, several reports have demonstrated proteolytic activity of CTSD for some substrates even in neutral pH environments [12, 37, 38]. In addition, interestingly, the CTSD precursor was recently reported to function as phosphatase in neutral pH environments [39]. These findings may provide a rationale for mitochondrial localization of CTSD, and concurrently, may suggest novel functions of CTSD in mitochondria. Further analysis is needed.

NCL is characterized by accumulation of ceroid lipofuscin containing the mitochondrial F1F0 ATPase [1, 2]. Involvement of autophagy has been demonstrated experimentally [40, 41], and the pathophysiology of NCL is explained, in part, by incomplete digestion of mitochondrial products by autophagy [42, 43]. Therefore, NCLs are caused by lysosomal disorders [1]. Among them, patients with CTSD dysfunction (CLN10), manifest early onset in NCL [1]. If mitochondrial forms of CTSD have a regulatory role in mitochondrial functions, the most severe forms of the disease may be attributed to the dysfunction of both mitochondrial and lysosomal CTSDs. In addition, some double-membrane structures in CLN10 containing mitochondrial F1F0 ATPase may reflect not only autophagic vacuoles (AVs), but also abnormal mitochondria themselves. Consistent with this idea, Sulzer et al. listed mitochondria as the only candidate, other than AVs [42], and some autolysosome-like bodies in CLN10 models [2] [40] sharing a common appearance with onion-like mitochondria, which are an abnormal form of mitochondria [44–47].

So far, CTSD has always been viewed as a lysosomal protein. The existence of mitochondrial CTSD demonstrated here may provide many new insights and may suggest new research questions regarding classical conceptions about CTSD.

## Experimental Procedures

### Cell lines

The following cell lines were purchased from the American Type Culture Collection: HeLa (tissue, cervical cancer, female), SK-BR-3 (tissue, breast cancer, female), HepG2 (tissue, liver cancer, male), and 293 (tissue, embryonic kidney). Cells were cultured in DMEM (Sigma). All media were supplemented with 10% fetal bovine serum. Cells were maintained at 37°C in a humidified chamber with 5% CO_2_.

### Establishment of KD cell lines

We established a CTSD-KD cell line using HeLa cells, as previously described [16]. CTSD expression was inhibited in this cell line by retroviral expression of short-hairpin RNA (shRNA) against the CTSD sequence [18]. We also established HeLa-cont cells using an empty retroviral vector. All oligonucleotides are listed in Supplementary Table S1.

### Plasmid construction

For construction of plasmids containing C-terminal enhanced green fluorescent protein-CTSD (pEGFP-CTSD), the nucleotide sequence of CTSD was PCR-amplified using the primers, CTSD-F1 and CTSD-R1. PCR products were digested with Nhe I and Hind III and ligated into pC-EGFP [19] cut with the same enzymes.

For construction of plasmids containing CTSD (pCTSD), pEGFP-CTSD was subjected to inverse PCR using the primers, CTSD-F2 and CTSD-R2, and the product was self-ligated using a KOD-Plus-Mutagenesis Kit (TOYOBO).

For construction of plasmids containing deletion mutants of C-terminal EGFP-CTSD and CTSD (Δ2-20, Δ2-35, Δ2-64, and Δ21-35), pEGFP-CTSD and pCTSD were subjected to inverse PCR using the primers, Δ-20F and Δ2-R, Δ-35F and Δ2-R, Δ-64F and Δ2-R, and Δ-35F and Δ21-R, respectively, and the product was self-ligated using a KOD-Plus-Mutagenesis Kit (TOYOBO). All primers are listed in Supplementary Table S1.

### Recombinant adenovirus construction

Replication-deficient recombinant viral Ad-CTSD was generated from pCTSD and purified as described previously [20]. Briefly, DNA fragments obtained by restriction of each plasmid vector were blunted using T4 DNA polymerase, ligated into the SmiI site of the cosmid, pAxCAwtit (Takara), which contains the CAG promoter and the entire genome of type 5 adenovirus, except the E1 and E3 regions. Recombinant adenoviruses were generated using *in vitro* homologous recombination in the 293 cell line with the cDNA-inserted pAxCAwtit and the adenovirus DNA terminal–protein complex. Viruses were propagated in the 293 cell line and purified with two rounds of CsCl density centrifugation. Viral titers were determined with a limiting dilution bioassay using 293 cells.

### Transfection

Cells seeded in 35-mm glass bottom dishes, 8-chamber slides (0.7 cm^2^ culture area), or 100-mm dishes were transfected with plasmids (2 μg /dish, 0.4 μg /well, or 10 μg /dish respectively) using FuGENE6 transfection reagent (Promega), according to the manufacturer’s instructions.

### Adenoviral infection

For transfection, cells were seeded (2×10^6^ cells/dish) in 100-mm dishes. Viral solutions were added to these cell monolayers, and cells were incubated at 37°C for 120 min with brief agitation every 20 min. This was followed by addition of culture medium, and cells were returned to the 37°C incubator.

### Amino acid sequence analyses of CTSD

The amino acid sequence of CTSD was analyzed using bioinformatics tools, DeepLoc-1.0 [21], DeepLoc-2.0 [22], MitoFates [23], and MITOPROT II [24].

### Analyses of confocal microscopy images

Confocal microscopy images were taken with a FLUOVIEW FV3000 confocal laser scanning microscope (Olympus). Line-scan profiles were acquired using MetaMorph ver. 7.8 (Molecular Devices).

### Proteinase K Protection Assay

HeLa and 293 cells (4×10^6^ cells/100-mm dish) were cultured under normal conditions for 24 h. HeLa cells (2×10^6^ cells/100-mm dish) 24 h after transfection with pEGFP-CTSD, or HeLa cells (2×10^6^ cells/100-mm dish) 24 h after infection with Ad-CTSD were harvested and suspended in 10 mM HEPES-KOH pH 7.5, 220 mM mannitol, 70mM sucrose. Suspended cells were homogenized using a syringe with a 27-gauge needle. Cell homogenates were centrifuged at 300 × g at 4 °C for 5 min to remove pellets containing unbroken cells and nuclei. Supernatants were centrifuged at 7,000 × g at 4 °C for 10 min to obtain pellets of crude mitochondria. Pellets of crude mitochondria were resuspended in 10 mM HEPES-KOH pH 7.5, 220 mM mannitol, 70 mM sucrose, and incubated with 0, 100, 200, 400 μg/mL of proteinase K, or 400 μg/mL of proteinase K with 1% Triton X-100 at 37 °C for 30 min. Subsequently, 1 mM phenylmethylsulfonyl fluoride (PMSF) was added to stop the reaction, followed by a 20 min incubation on ice. Samples were subjected to Western blotting.

### Sucrose Gradient

HeLa cells (1.2×10^7^ cells/100-mm dish) cultured under normal conditions for 24 h were harvested and suspended in 10 mM HEPES-KOH pH 7.5, 0.3 M sucrose containing Complete Protease Inhibitor Cocktail (Roche). Suspended cells were homogenized using a syringe with a 27-gauge needle. The cell homogenate was centrifuged at 300 × g at 4 °C for 5 min to remove unbroken cells and nuclei. Supernatant (1 vol.) was layered on top of a sucrose gradient consisting of 1.8 M, 1.5M, 1.2 M, 0.9 M, 0.6 M sucrose in 10 mM HEPES-KOH pH 7.5, 5 vol. containing Complete Protease Inhibitor Cocktail (Roche). After centrifugation at 100,000 × g at 4 °C for 16 h, 12 fractions were collected from the top.

### Western blotting analysis

Proteins separated with SDS-PAGE were transferred onto polyvinylidene difluoride (PVDF) membranes (Millipore). Membranes were blocked with 5% skim milk in Tris-buffered saline with 0.05% Tween 20 for 1 h. Membranes were incubated with primary antibodies at 4 °C overnight, and subsequently with secondary antibodies for 1 h. Antibodies used were as follows: anti-CTSD mouse monoclonal (1:1000, CTD-19, NOVUS), anti-CTSD goat polyclonal (1:250, sc-6486, Santa Cruz Biotechnology), anti-TOMM20 (1:250, sc-17764, Santa Cruz Biotechnology), anti-ATP5F1B (1:400, PAB28448, Abnova), anti-SEC22B (1:200, sc-101267, Santa Cruz Biotechnology), anti-Golgin97 (1:200, sc-59820, Santa Cruz Biotechnology), anti-beta-actin (1:5000, A5316, Sigma-Aldrich), and anti-GFP (sc-9996, Santa Cruz Biotechnology), anti-rabbit IgG-HRP (1:5000, sc-2004, Santa Cruz Biotechnology), anti-mouse IgG-HRP (1:5000, sc-2005, Santa Cruz Biotechnology), and anti-goat IgG-HRP (1:5000, sc-2020, Santa Cruz Biotechnology)

### Analysis of CTSD localization in fixed cells

HeLa cells transfected with pEGFP-CTSD were subjected to none, one, pairs, or all of the following procedures for immunofluorescence (IF), 24 h after transfection. i) Fixation with 4% paraformaldehyde for 15 min. ii) Permeabilization with 0.1% Triton X-100 for 5 min. iii) Subsequent procedures for antigen-antibody reaction: washing 3x with phosphate-buffered saline (PBS) at room temperature, blocking with 3% bovine serum albumin (BSA) in PBS for 1 h, incubating with primary antibody, anti-TOMM20 (1:100, sc-17764, Santa Cruz Biotechnology) or anti-cytochrome *c* (1:100, 6H2.B4, Imgenex) in 3% BSA in PBS for 2 h at room temperature, washing 3x with PBS, incubating with secondary antibody, Alexa Fluor 546 (1:200, A-11003, Thermo Fisher Scientific) in 3% BSA in PBS for 1 h at room temperature, and washing 3x with PBS. Cells were mounted with VECTASHIELD H-1000 (Vector Laboratories) and observed using a FLUOVIEW FV3000 confocal laser scanning microscope (Olympus).

### Statistical analysis

Statistical analyses were performed in JMP 14.2.0 (SAS). Levels of significance were assessed using Student’s two-tailed t-tests. p < 0.05 was considered statistically significant.

## Data availability

All data are contained within this manuscript.

## Supporting information

This article contains supporting information.

## Acknowledgments

We thank T. Shibata, Y. Nakamura, K. Honjo, and Y. Sagami for their assistance.

## Author contribution

N.I. and H.A. conceptualization; N.I. and H.A. data curation; N.I. and H.A. funding acquisition; N.I. formal analysis; N.I. and H.A. investigation; N.I. methodology; H.A. project administration; H.A. supervision; N.I. and H.A. writing – original draft.

## Funding and additional information

This work was supported in part by grants from JSPS KAKENHI Grant Number JP-19K07703 (to NI), The National Cancer Center Research and Development Fund (29-E-1, 30-A-3) (to HA), JSPS KAKENHI Grant Numbers JP-19K07655, JP-17H06267, JP-20K20305, 22H02908, and 22K19412 (to HA), and The Naito Foundation (to HA).

## Conflict of interest

The authors declare no competing interests.

---

**Fig. S1:**
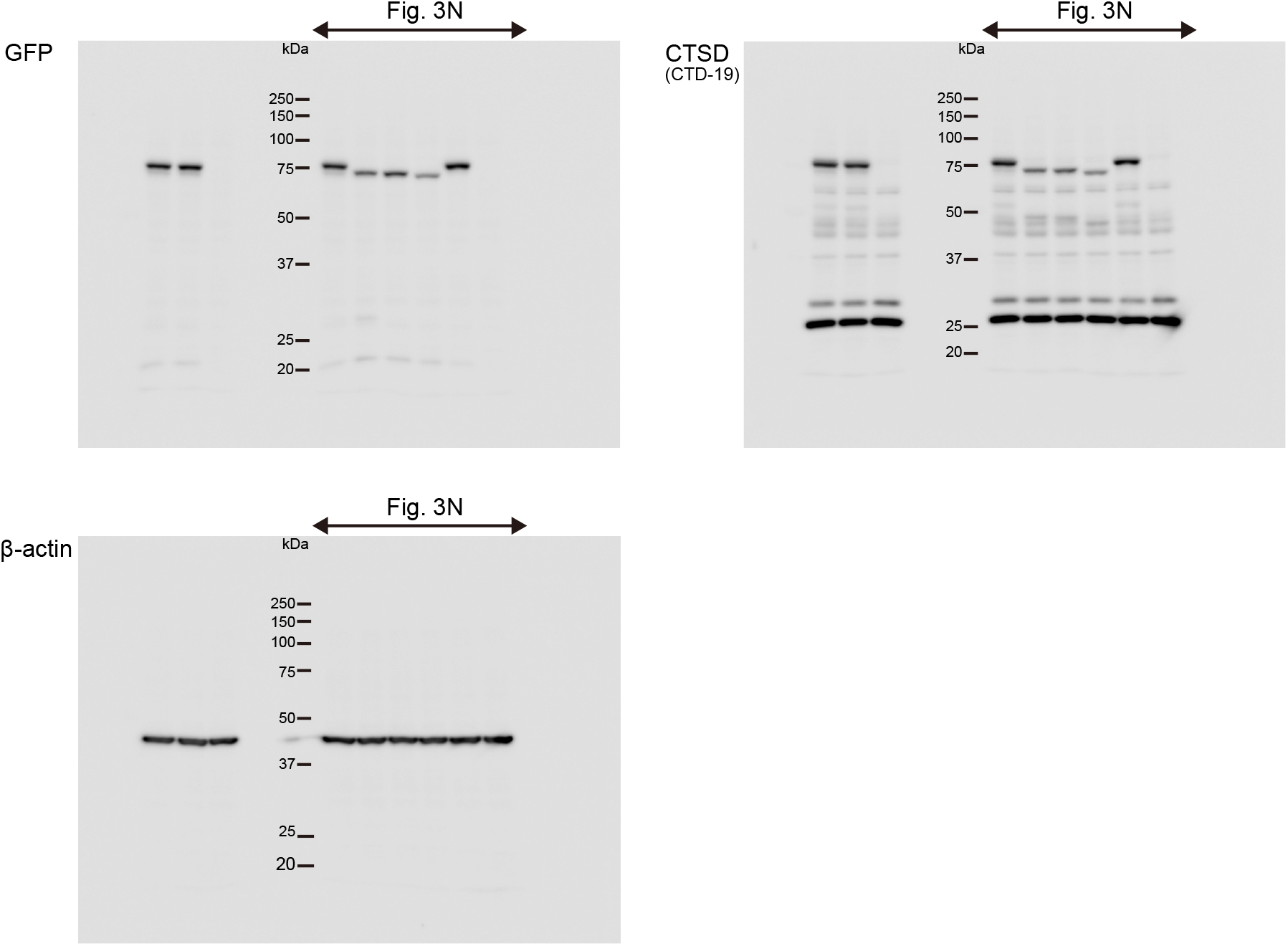
Source data of western blot in Fig. 3N

**Fig. S2:**
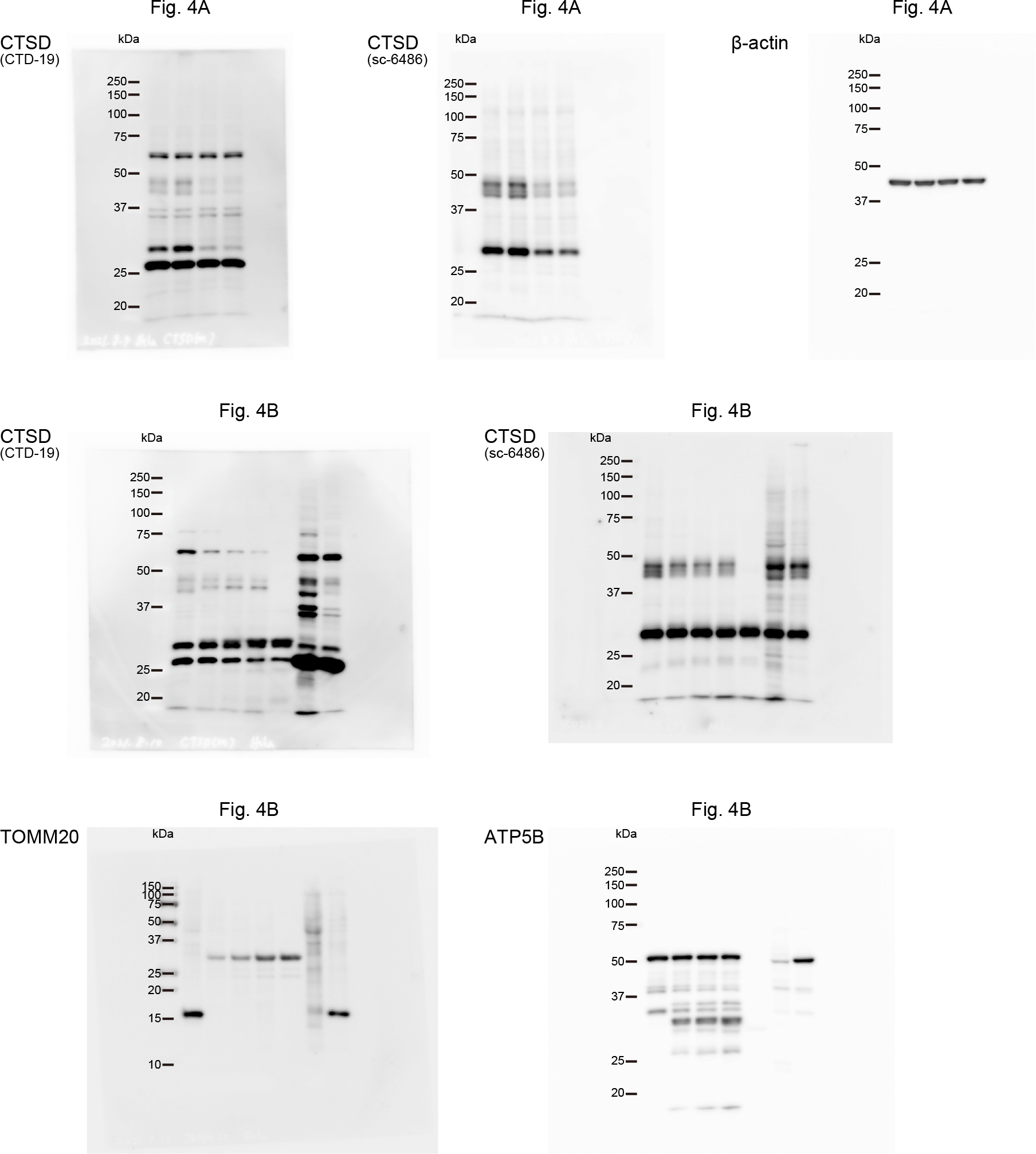
Source data of western blot in Fig. 4A and 4B

**Fig. S3:**
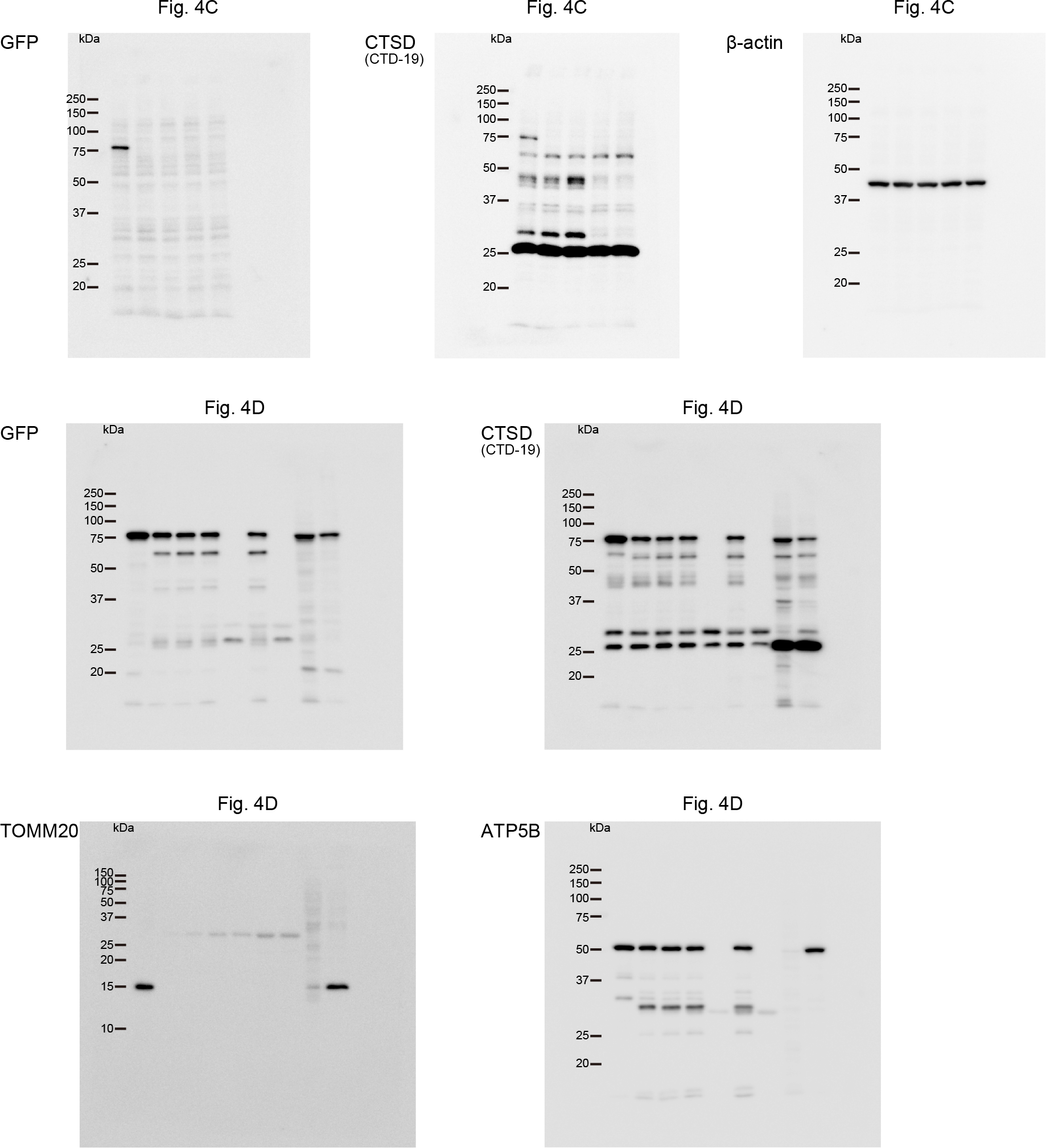
Source data of western blot in Fig. 4C and 4D

**Fig. S4:**
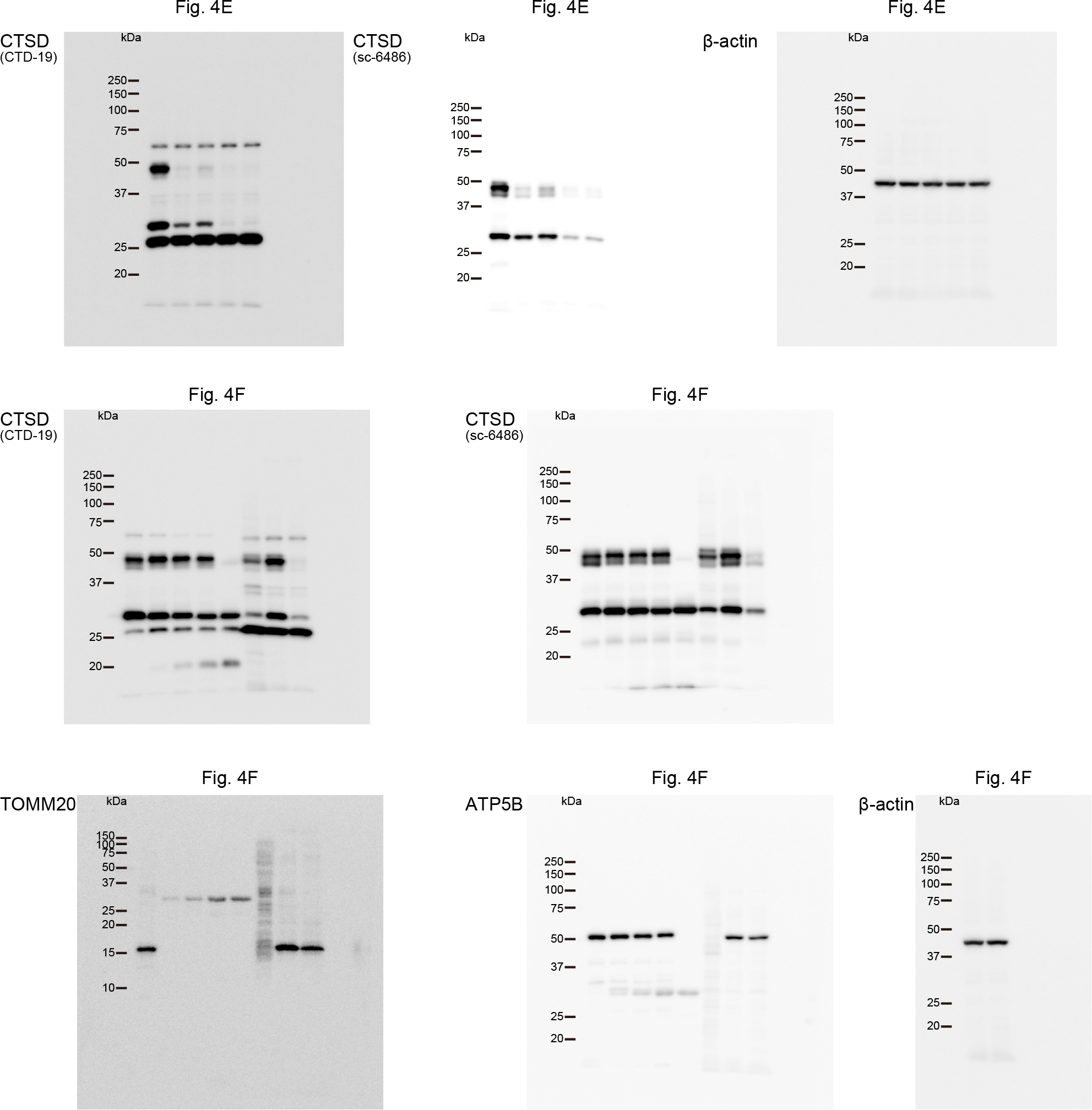
Source data of western blot in Fig. 4E and 4F

**Fig. S5:**
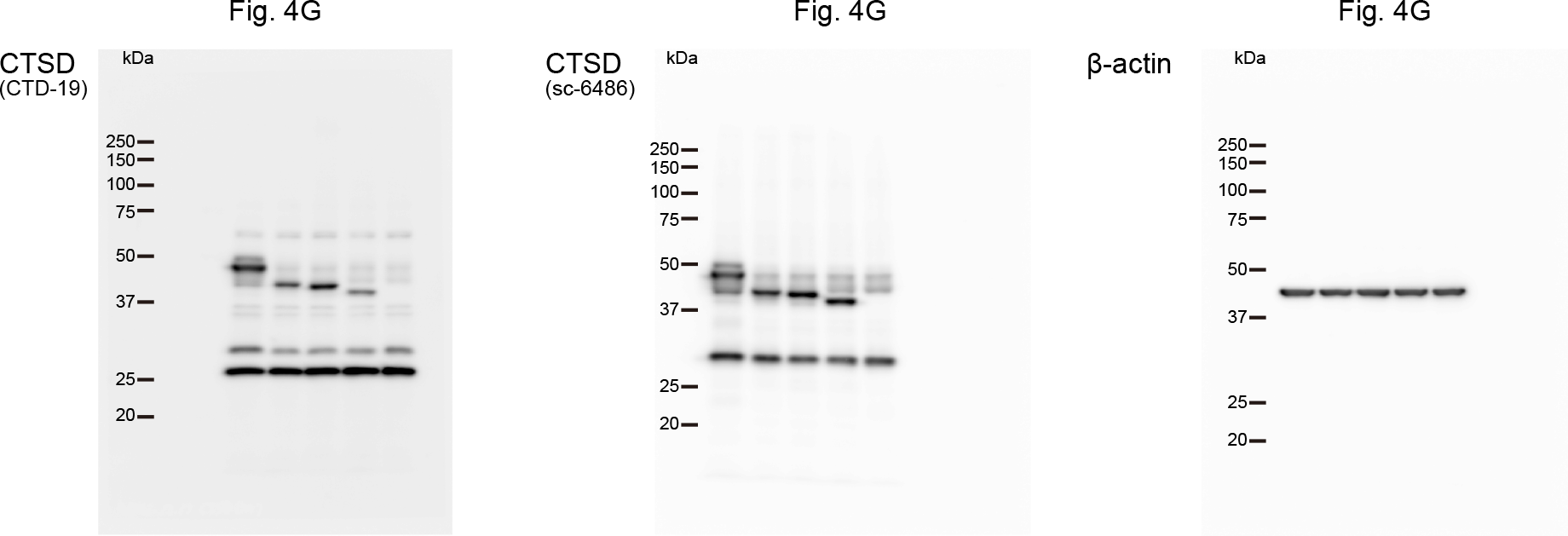
Source data of western blot in Fig. 4G

**Fig. S6:**
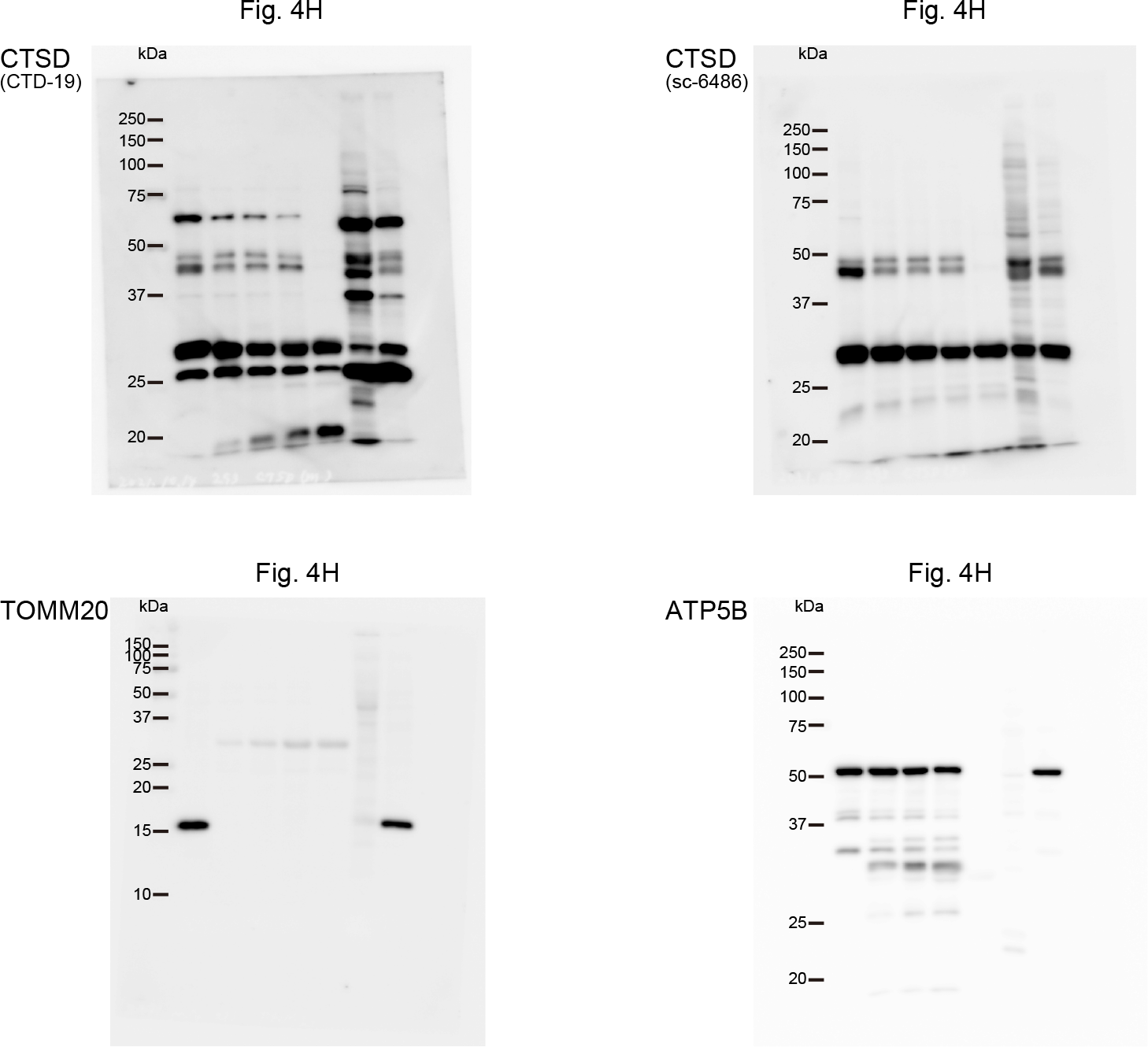
Source data of western blot in Fig. 4H

**Fig. S7:**
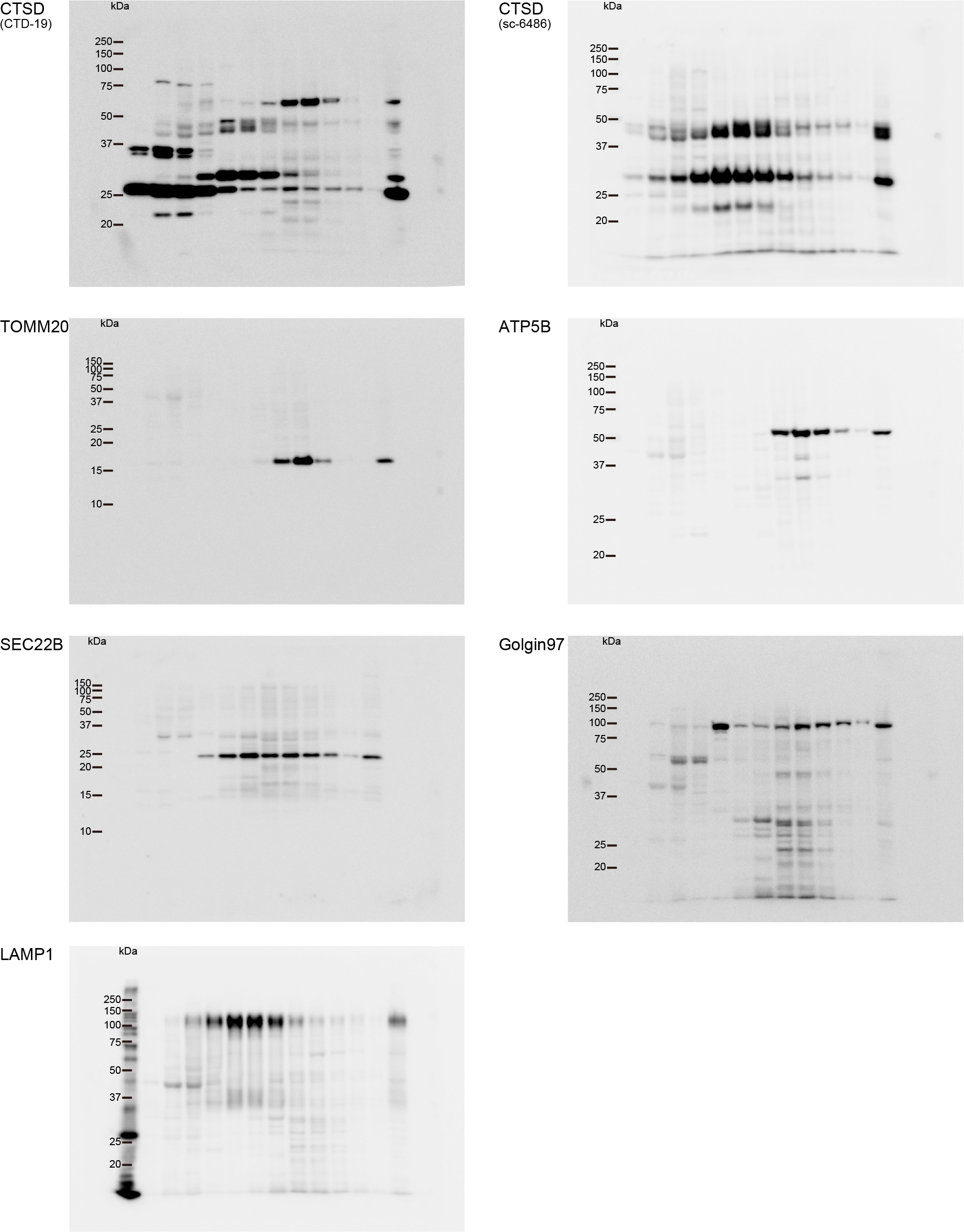
Source data of western blot in Fig. 4I

**Table S1.**
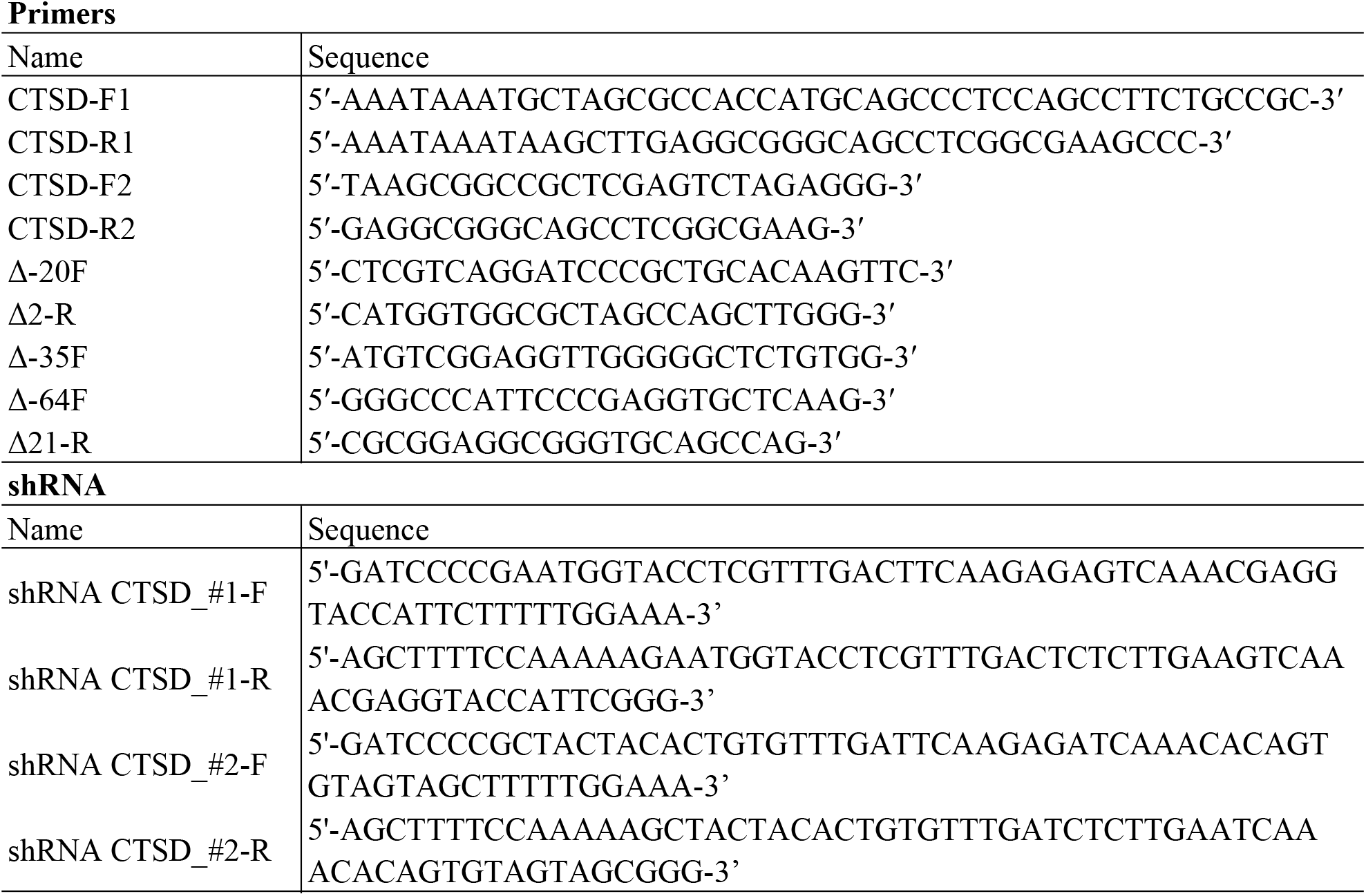
Oligonucleotides used in this paper.

**Table S2.**
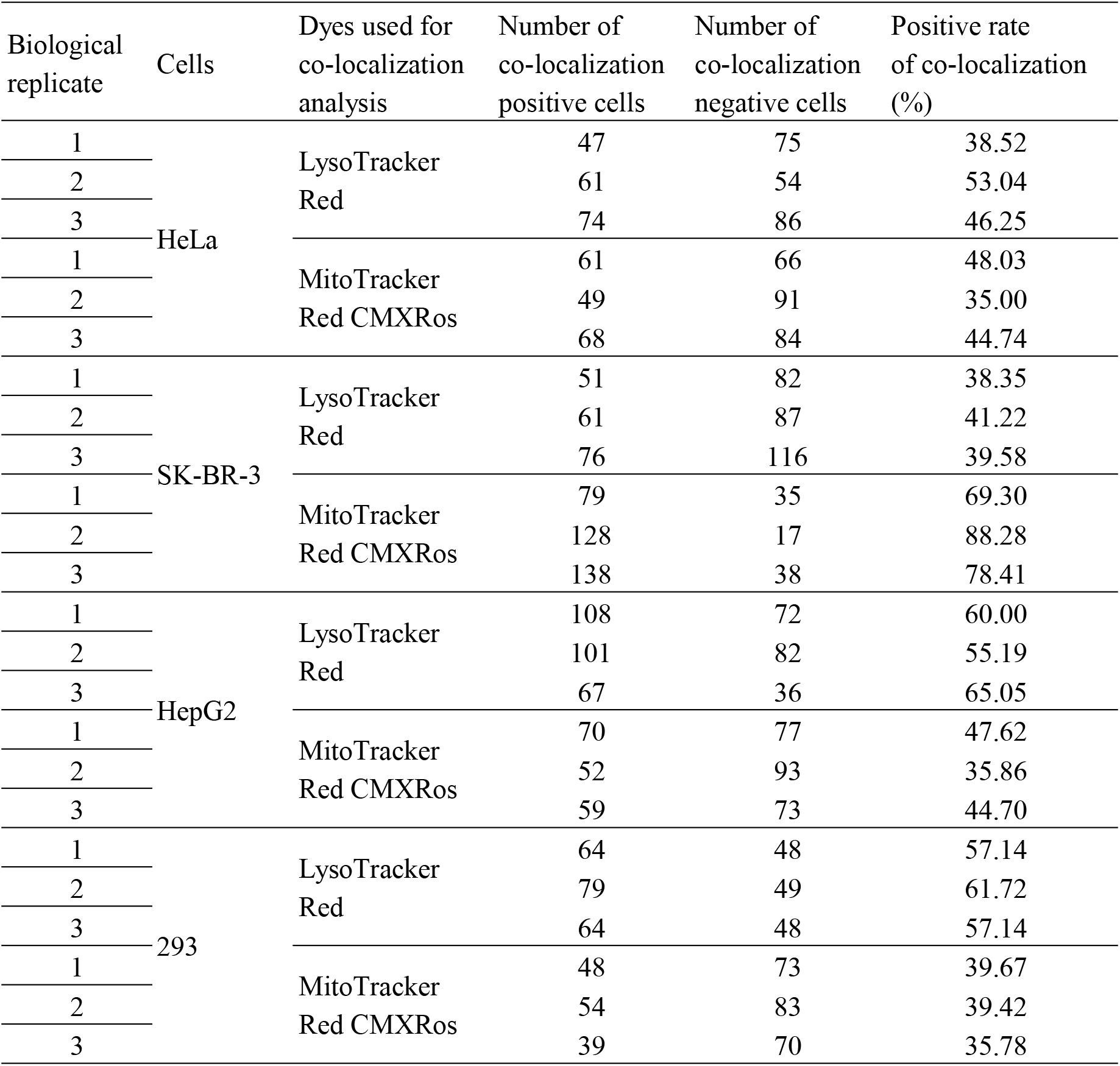
Subcellular localization of EGFP-CTSD expressed in various cell lines.

**Table S3.**
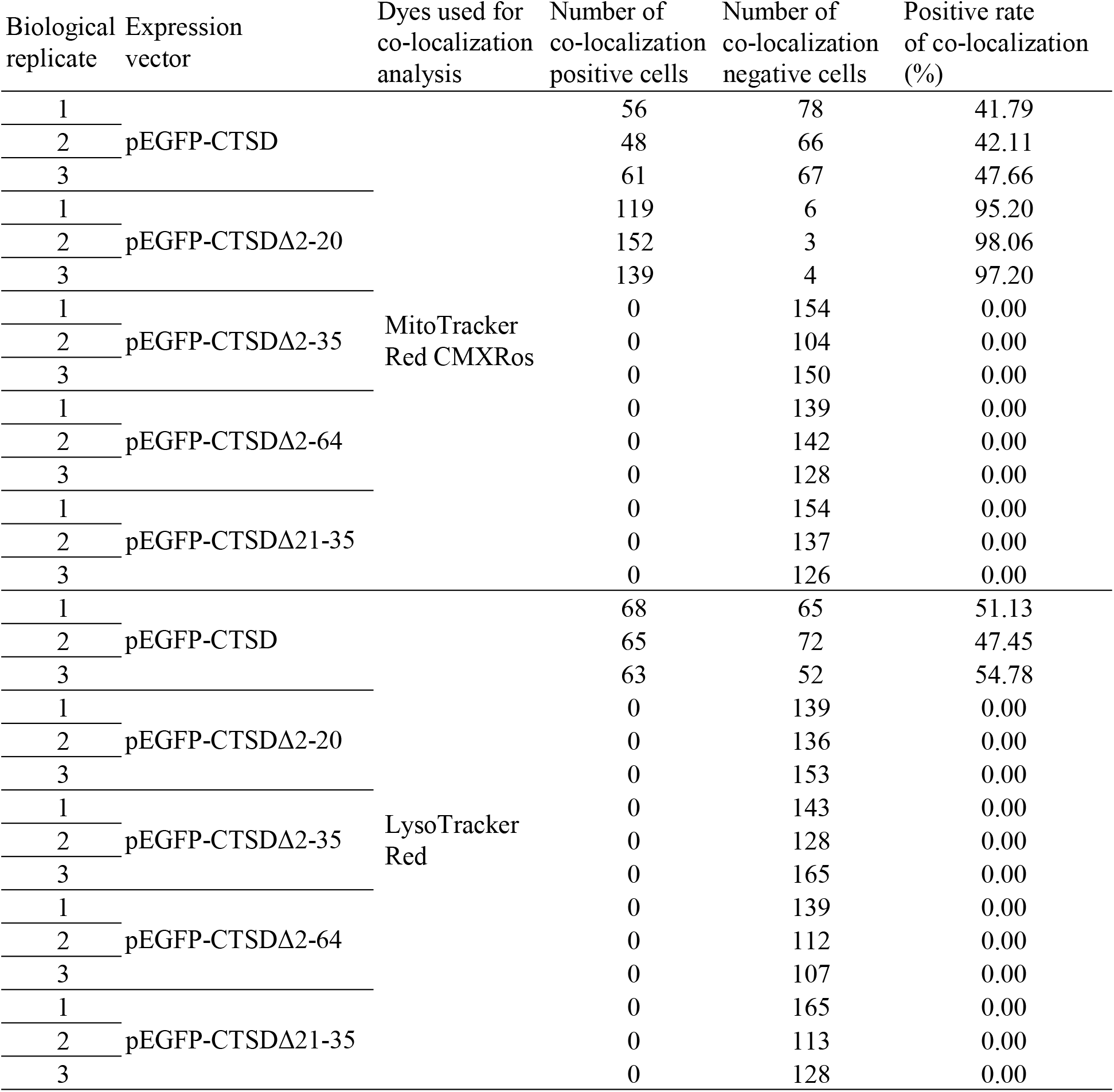
Subcellular localization of EGFP-CTSD and the deletion mutants expressed in HeLa cells.

## Notes

### Competing Interest Statement

The authors have declared no competing interest.

