## Supplementary figures for "Identification of a mitochondrial targeting sequence in cathepsin D and its localization in mitochondria"

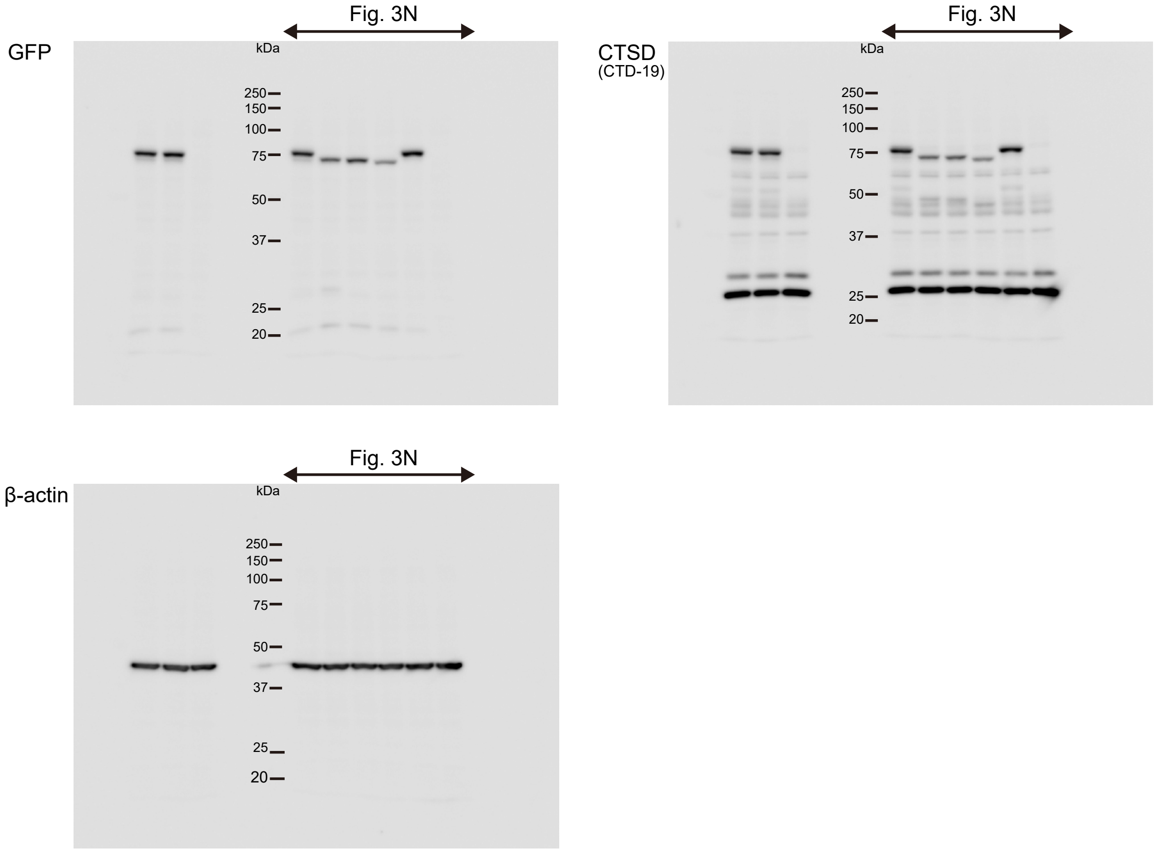


Fig. S1: Source data of western blot in Fig. 3N

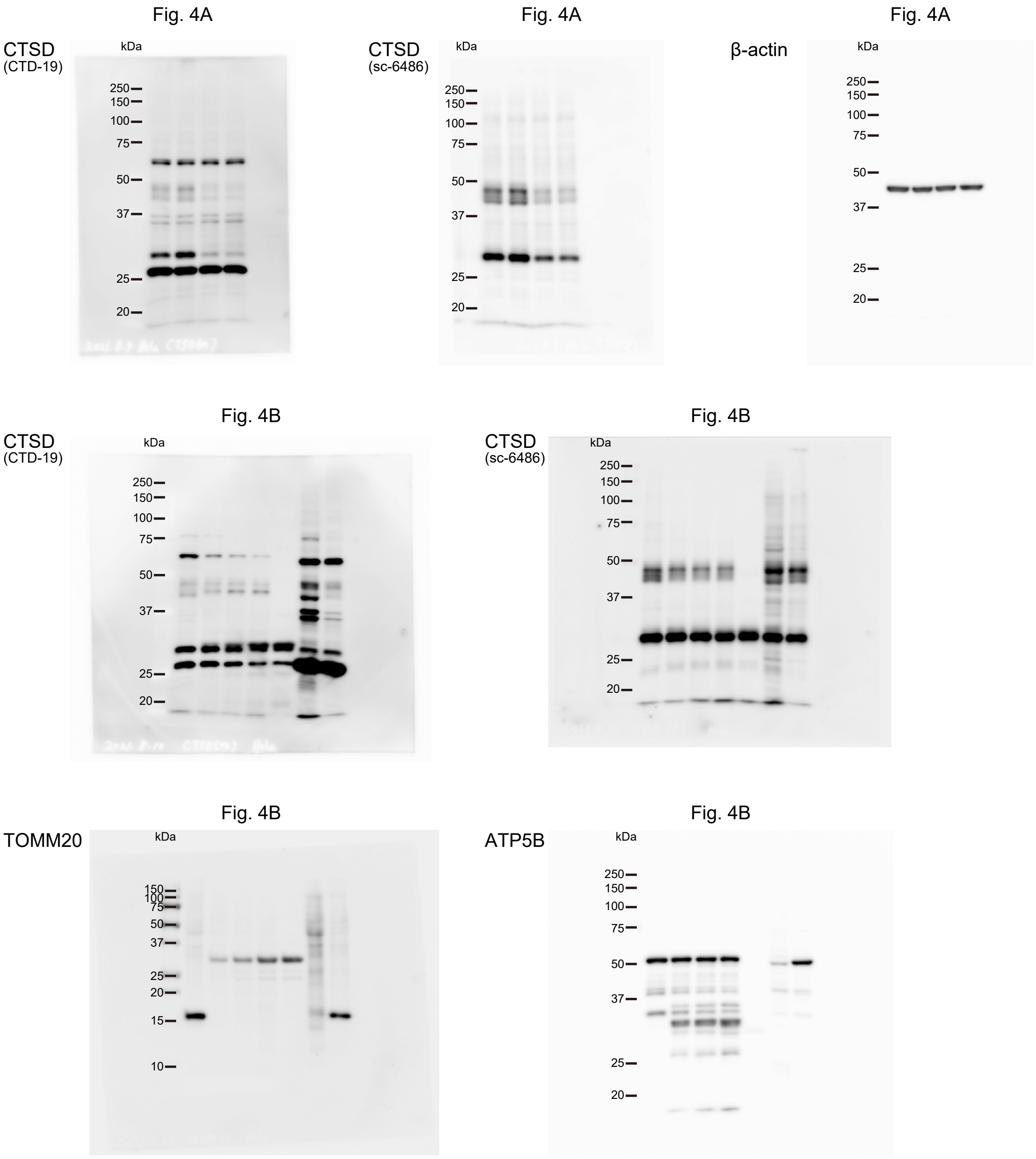


Fig. S2: Source data of western blot in Fig. 4A and 4B

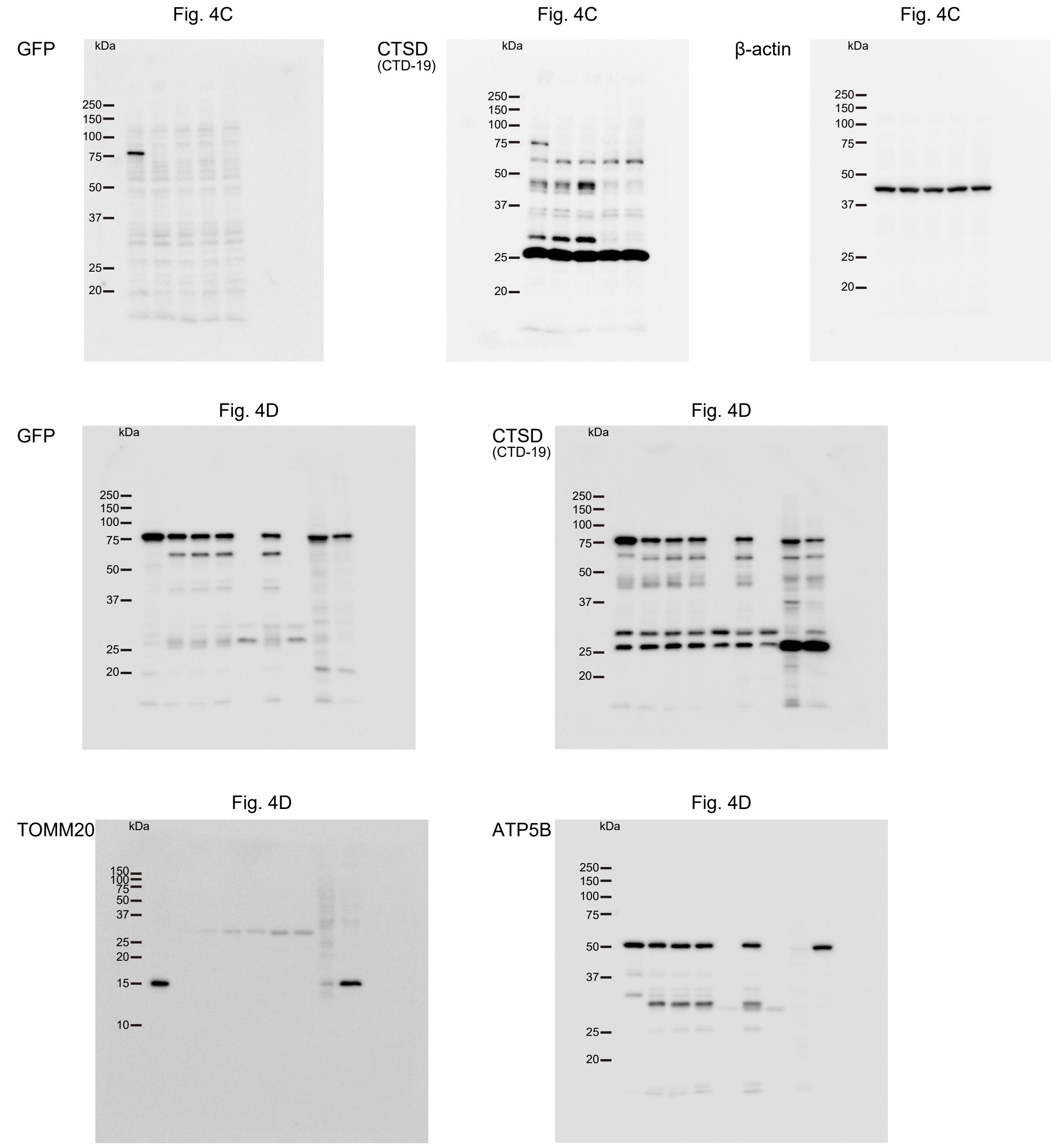


Fig. S3: Source data of western blot in Fig. 4C and 4D

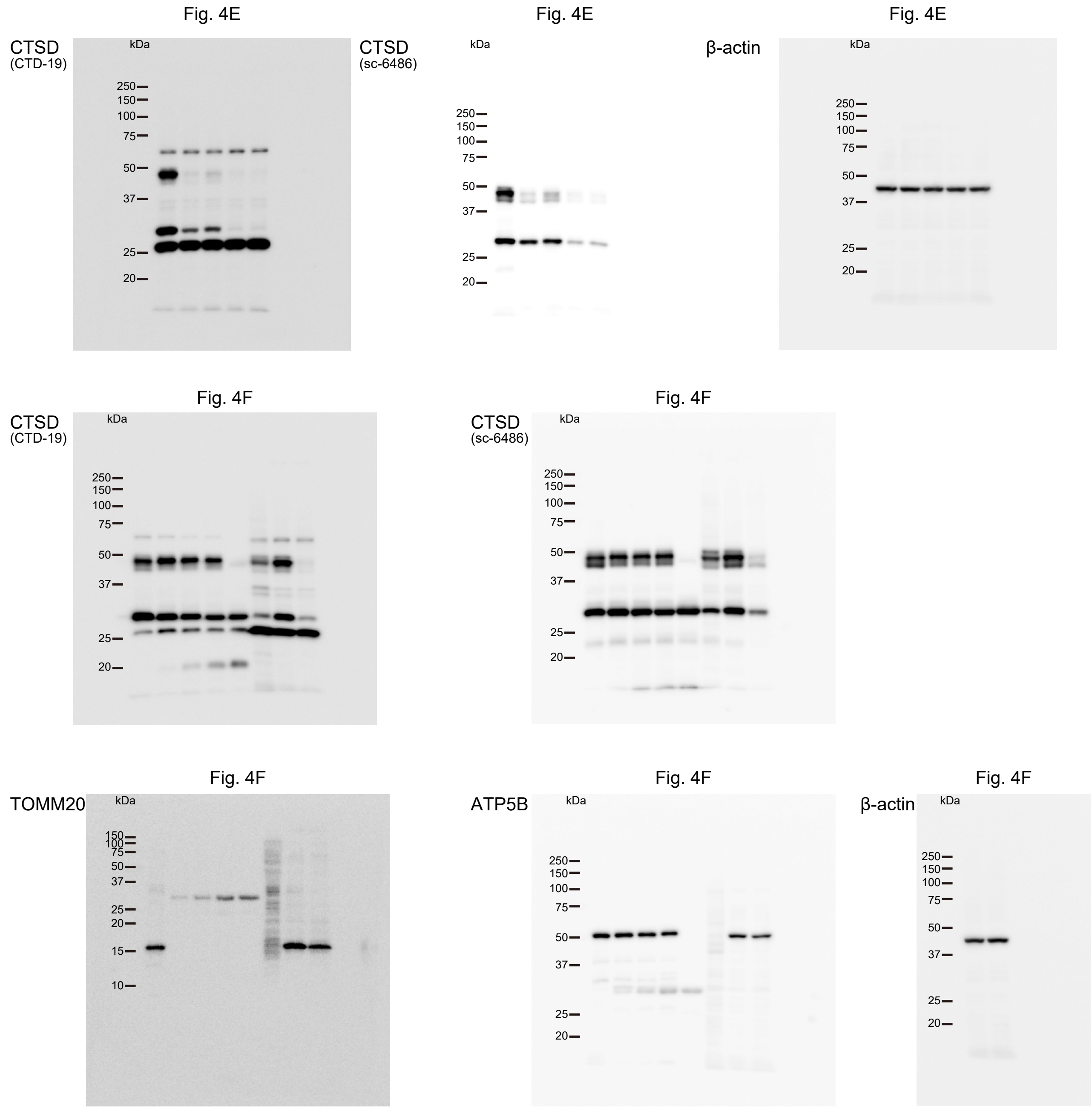


Fig. S4: Source data of western blot in Fig. 4E and 4F

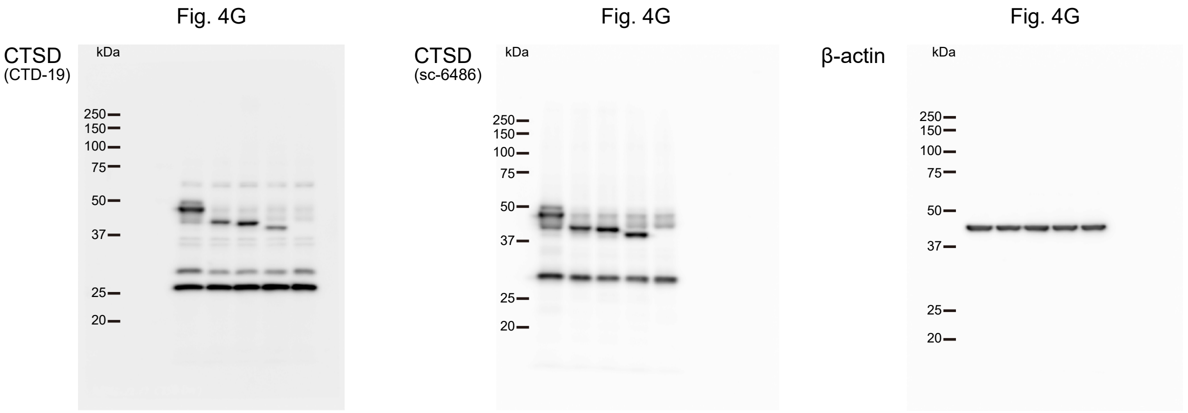


Fig. S5: Source data of western blot in Fig. 4G

Fig. 4H

CTSD
(CTD-19)

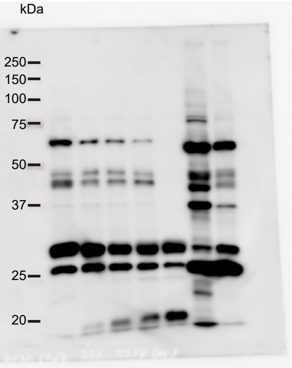


Fig. 4H

CTSD
(sc-6486)

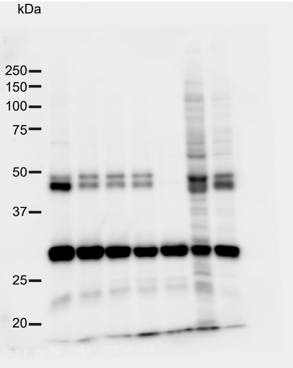


Fig. 4H

TOMM20

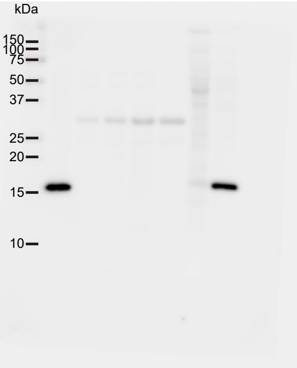


Fig. 4H

ATP5B

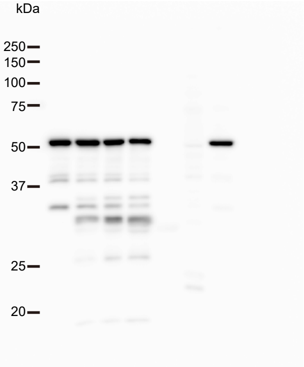


Fig. S6: Source data of western blot in Fig. 4H

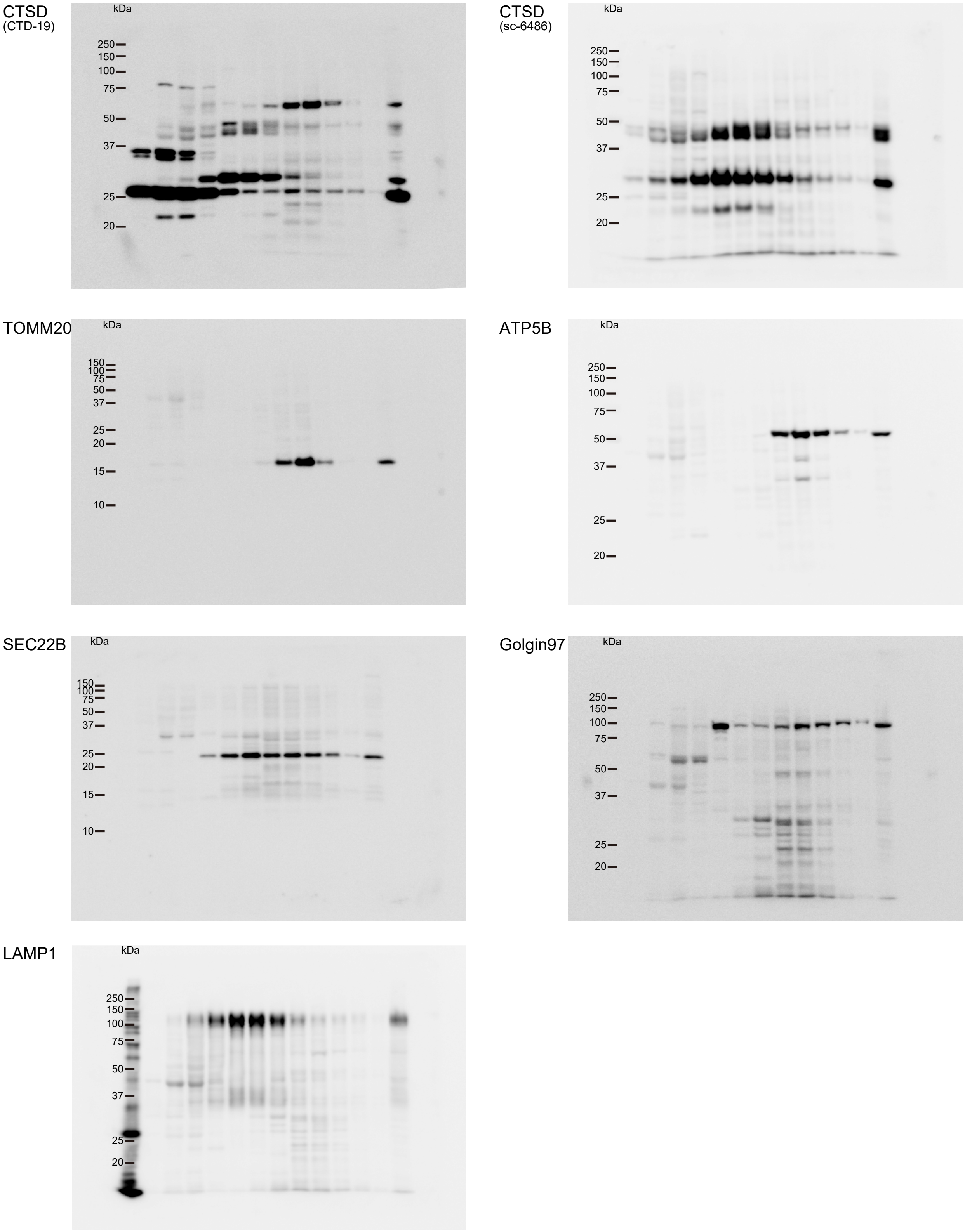


Fig. S7: Source data of western blot in Fig. 4I
