## Supplementary tables for "Identification of a mitochondrial targeting sequence in cathepsin D and its localization in mitochondria"

**Table S1. Oligonucleotides used in this paper**

**Primers**

| Name | Sequence |
|---|---|
| CTSD-F1 | 5′-AAATAAATGCTAGCGCCACCATGCAGCCCTCCAGCCTTCTGCCGC-3′ |
| CTSD-R1 | 5′-AAATAAATAAGCTTGAGGCGGGCAGCCTCGGCGAAGCCC-3′ |
| CTSD-F2 | 5′-TAAGCGGCCGCTCGAGTCTAGAGGG-3′ |
| CTSD-R2 | 5′-GAGGCGGGCAGCCTCGGCGAAG-3′ |
| Δ-20F | 5′-CTCGTCAGGATCCCGCTGCACAAGTTC-3′ |
| Δ2-R | 5′-CATGGTGGCGCTAGCCAGCTTGGG-3′ |
| Δ-35F | 5′-ATGTCGGAGGTTGGGGGCTCTGTGG-3′ |
| Δ-64F | 5′-GGGCCCATTCCCGAGGTGCTCAAG-3′ |
| Δ21-R | 5′-CGCGGAGGCGGGTGCAGCCAG-3′ |

**shRNA**

| Name | Sequence |
|---|---|
| shRNA CTSD_#1-F | 5'-GATCCCCGAATGGTACCTCGTTTGACTTCAAGAGAGTCAAACGAGG TACCATTCTTTTTGGAAA-3’ |
| shRNA CTSD_#1-R | 5'-AGCTTTTCCAAAAAGAATGGTACCTCGTTTGACTCTCTTGAAGTCAA ACGAGGTACCATTCGGG-3’ |
| shRNA CTSD_#2-F | 5'-GATCCCCGCTACTACACTGTGTTTGATTCAAGAGATCAAACACAGT GTAGTAGCTTTTTGGAAA-3’ |
| shRNA CTSD_#2-R | 5'-AGCTTTTCCAAAAAGCTACTACACTGTGTTTGATCTCTTGAATCAA ACACAGTGTAGTAGCGGG-3’ |

**Table S2. Subcellular localization of EGFP-CTSD expressed in various cell lines.**

| Biological replicate | Cells | Dyes used for co-localization analysis | Number of co-localization positive cells | Number of co-localization negative cells | Positive rate of co-localization (%) |
|---|---|---|---|---|---|
| 1 | HeLa | LysoTracker Red | 47 | 75 | 38.52 |
| 2 | | | 61 | 54 | 53.04 |
| 3 | | | 74 | 86 | 46.25 |
| 1 | | MitoTracker Red CMXRos | 61 | 66 | 48.03 |
| 2 | | | 49 | 91 | 35.00 |
| 3 | | | 68 | 84 | 44.74 |
| 1 | SK-BR-3 | LysoTracker Red | 51 | 82 | 38.35 |
| 2 | | | 61 | 87 | 41.22 |
| 3 | | | 76 | 116 | 39.58 |
| 1 | | MitoTracker Red CMXRos | 79 | 35 | 69.30 |
| 2 | | | 128 | 17 | 88.28 |
| 3 | | | 138 | 38 | 78.41 |
| 1 | HepG2 | LysoTracker Red | 108 | 72 | 60.00 |
| 2 | | | 101 | 82 | 55.19 |
| 3 | | | 67 | 36 | 65.05 |
| 1 | | MitoTracker Red CMXRos | 70 | 77 | 47.62 |
| 2 | | | 52 | 93 | 35.86 |
| 3 | | | 59 | 73 | 44.70 |
| 1 | 293 | LysoTracker Red | 64 | 48 | 57.14 |
| 2 | | | 79 | 49 | 61.72 |
| 3 | | | 64 | 48 | 57.14 |
| 1 | | MitoTracker Red CMXRos | 48 | 73 | 39.67 |
| 2 | | | 54 | 83 | 39.42 |
| 3 | | | 39 | 70 | 35.78 |

**Table S3. Subcellular localization of EGFP-CTSD and the deletion mutants expressed in HeLa cells.**

| Biological replicate | Expression vector | Dyes used for co-localization analysis | Number of co-localization positive cells | Number of co-localization negative cells | Positive rate of co-localization (%) |
|---|---|---|---|---|---|
| 1 | pEGFP-CTSD | MitoTracker Red CMXRos | 56 | 78 | 41.79 |
| 2 | | | 48 | 66 | 42.11 |
| 3 | | | 61 | 67 | 47.66 |
| 1 | pEGFP-CTSDΔ2-20 | | 119 | 6 | 95.20 |
| 2 | | | 152 | 3 | 98.06 |
| 3 | | | 139 | 4 | 97.20 |
| 1 | pEGFP-CTSDΔ2-35 | | 0 | 154 | 0.00 |
| 2 | | | 0 | 104 | 0.00 |
| 3 | | | 0 | 150 | 0.00 |
| 1 | pEGFP-CTSDΔ2-64 | | 0 | 139 | 0.00 |
| 2 | | | 0 | 142 | 0.00 |
| 3 | | | 0 | 128 | 0.00 |
| 1 | pEGFP-CTSDΔ21-35 | | 0 | 154 | 0.00 |
| 2 | | | 0 | 137 | 0.00 |
| 3 | | | 0 | 126 | 0.00 |
| 1 | pEGFP-CTSD | LysoTracker Red | 68 | 65 | 51.13 |
| 2 | | | 65 | 72 | 47.45 |
| 3 | | | 63 | 52 | 54.78 |
| 1 | pEGFP-CTSDΔ2-20 | | 0 | 139 | 0.00 |
| 2 | | | 0 | 136 | 0.00 |
| 3 | | | 0 | 153 | 0.00 |
| 1 | pEGFP-CTSDΔ2-35 | | 0 | 143 | 0.00 |
| 2 | | | 0 | 128 | 0.00 |
| 3 | | | 0 | 165 | 0.00 |
| 1 | pEGFP-CTSDΔ2-64 | | 0 | 139 | 0.00 |
| 2 | | | 0 | 112 | 0.00 |
| 3 | | | 0 | 107 | 0.00 |
| 1 | pEGFP-CTSDΔ21-35 | | 0 | 165 | 0.00 |
| 2 | | | 0 | 113 | 0.00 |
| 3 | | | 0 | 128 | 0.00 |
